# Sub-cellular chemical mapping in bacteria using correlated cryogenic electron and mass spectrometry imaging

**DOI:** 10.1101/2025.08.29.673036

**Authors:** Hannah Ochner, Buse Isbilir, Sonja Blasche, David Scheidweiler, Yuexuan Zhang, Zhexin Wang, Tom Smith, Catarina Franco, Rob Bradley, Kiran R. Patil, Tanmay A.M. Bharat

**Affiliations:** Structural Studies Division, MRC Laboratory of Molecular Biology; Francis Crick Avenue, Cambridge CB2 0QH, United Kingdom; Cell Biology Division, MRC Laboratory of Molecular Biology; Francis Crick Avenue, Cambridge CB2 0QH, United Kingdom; MRC Toxicology Unit, University of Cambridge; Tennis Court Road, Cambridge CB2 1QR, United Kingdom

## Abstract

**Electron cryomicroscopy (cryo-EM) allows high-resolution spatial visualization of biological specimens, however, it is challenging to chemically identify densities observed in cryo-EM. To overcome this, we combined cryo-EM with chemical imaging using focused ion beam secondary ion mass spectrometry (FIB-SIMS) for integrated spatio-chemical analysis of untagged specimens. We show that our correlative workflow permits sub-cellular localisation of molecules inside bacterial cells and is compatible with cryogenic light microscopy and FIB-milled lamellae of multicellular specimens. To highlight biological insights enabled by the workflow, we studied the uptake of Bisphenol-AF, a widespread chemical pollutant, by environmental bacteria, revealing the storage of these chemicals within cytosolic phase-separated aggregates in pollutant-exposed cells, where they cannot be removed by the bacterial efflux machinery despite its robust upregulation. These mechanistic insights were directly facilitated by the versatile cryo-EM-FIB-SIMS technique, showing that it is an effective avenue to map elemental and molecular signatures in near-native biological samples, which can be extended in the future for multiple applications in cell biology and imaging.**

## Introduction

Electron cryomicroscopy (cryo-EM) provides high spatial resolution for visualizing a wide range of biological samples from purified molecules to cells and tissues^1–3^. However, there is no direct way of assigning chemical identity to the densities observed in electron micrographs, which means that different molecules present within the sample cannot be easily distinguished from each other, especially within the cellular context^1^. Mass spectrometry (MS), on the other hand, is a chemically specific technique, allowing direct chemical identification of the molecules in a sample and thus providing highly complementary information to cryo-EM structural data. While many MS techniques are performed on bulk samples, MS can also be conducted in an imaging modality, which allows simultaneous detection of the spatial distributions of different molecules in a specimen^4–8^. In particular, secondary ion mass spectrometry (SIMS), utilizing a range of different ion beams for the pixel-by-pixel ionization of the specimen in conjunction with extraction of the resulting secondary ions into mass analyzers such as time-of-flight (ToF-SIMS) and Orbitrap (OrbiSIMS) mass spectrometers, has been shown to be a versatile tool for studying the chemical composition of biological specimens^9–15^. While SIMS applications range from single cell to tissue imaging, and from elemental ion detection to metabolomics and proteomics^9–14,16^, the attainable spatial resolution is limited by the size of the ion beam^4^. To detect the presence or absence of molecules of interest (or their fragments) along with their sub-cellular localization, single-cell measurements of untagged specimens with both high spatial resolution and chemical sensitivity are required. Examples of biological processes that would benefit from such imaging include, but are not limited to, drug uptake by cells, pollutant uptake by bacteria and symbiotic metabolite sharing within microbiomes. Thus, combining the capabilities of cryo-EM and imaging MS would extend the capabilities of both these techniques, providing images with unprecedented explanatory power for mechanistic structural and cellular studies of a wide range of biological processes.

To this end, we have developed a workflow to perform correlated cryo-EM with MS imaging of vitrified biological specimens using a focused ion beam (FIB)-SIMS instrument at cryogenic temperatures. We provide proof-of-principle data for tagged and untagged biological specimens and extend our approach to show that the technique is compatible with cryo-light microscopy (cryo-LM), as well as with multicellular, tissue-like specimens, which can be imaged in the form of FIB-milled lamellae. These examples demonstrate the versatility of the spatio-chemical imaging workflow, which can be tailored to the specific requirements of the biological specimen. To highlight the utility of our integrated workflow, we studied bioaccumulation of environmental contaminants in bacterial cells. Specifically, we studied the uptake of a widespread fluorinated pollutant, Bisphenol-AF (BPAF), by environmental bacteria, showing that these molecules are concentrated in the cytosol inside phase-separated aggregates and cannot be removed by the drug efflux machinery despite its upregulation by the bacteria.

Our results on BPAF bioaccumulation underscore the complementarity of cryo-EM and MS, showing how effectively combining the two imaging modalities supports biological discovery.

## Results

### Integrated cryo-EM and FIB-SIMS imaging workflow

We developed a correlative workflow enabling chemical mapping at the sub-cellular scale in bacteria (Fig. 1a) under cryogenic conditions in vitrified specimens (Methods). First, two-dimensional (2D) cryo-EM images of the sample are collected (Fig. S1a), followed by specimen transfer to the cryo-FIB-SIMS instrument in which MS imaging is performed on the areas pre-imaged by cryo-EM to allow correlation of the cryo-EM and cryo-FIB-SIMS data sets. During specimen transfer, the stability of the sample is enhanced by applying an organometallic coating to the backside of the sample, leading to an improved MS signal as sample movement and inhomogeneities of the sample support are reduced (Fig. S2).

**Figure 1:**
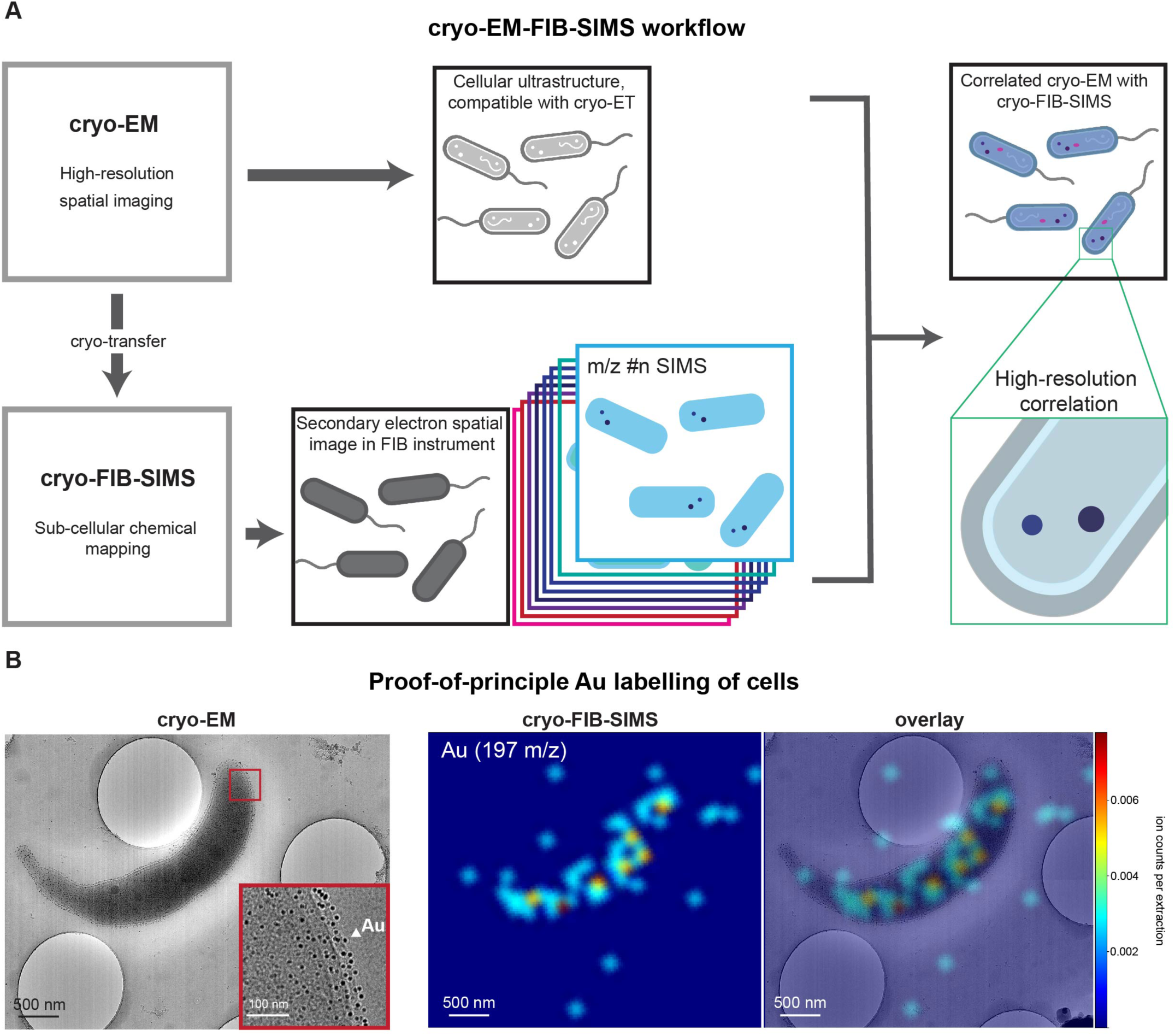
Correlative cryo-EM-FIB-SIMS workflow. **(A)** Schematic of the cryo-EM-FIB-SIMS workflow. As a first step, vitrified specimens are imaged by cryo-EM, yielding high spatial resolution images detailing the cellular ultrastructure. Subsequently, the samples are transferred to the cryo-FIB-SIMS instrument where imaging MS is performed on the sample regions pre-imaged by cryo-EM. In addition to the chemical information from SIMS imaging, secondary electron FIB images are also collected which are useful for correlation of the cryo-FIB-SIMS and the cryo-EM data. The data from both imaging modalities can then be overlaid, connecting the chemical information from cryo-FIB-SIMS with the high spatial resolution information from cryo-EM. **(B)** Proof-of-principle example of the workflow using gold nanoparticle-labelled *C. crescentus* cells. The S-layer of the cells has been labelled with gold nanoparticles resulting in a coating of the entire cell with gold as demonstrated by cryo-EM (left, gold nanoparticles appear as black dots around the cell). The inset is a high-magnification image of the area highlighted by the red box, gold nanoparticles are indicated by a white arrowhead. This is confirmed by the cryo-FIB-SIMS data which shows that the gold (Au, 197 m/z) signal co-localizes with the cell and is distributed along its length (right). The colour scale of the MS images encodes the detected ion count per extraction. Discrepancies between the cryo-FIB-SIMS gold signal and the distribution of gold nanoparticles in the cryo-EM image are likely the result of the difference in resolution between the two imaging modalities as well as nonuniform ionization yields across the cell.

The cryo-FIB-SIMS instrument is a focused ion beam scanning electron microscope (FIB-SEM) equipped with a time-of-flight (ToF) mass spectrometer. To perform SIMS imaging, a focused gallium (Ga) ion beam, with an unbinned pixel size range in MS imaging mode of approximately 10-30 nm, is used to scan the field of view, ablating material by interacting with the specimen (Fig. S1b). The secondary ions produced by this interaction are extracted into the ToF mass spectrometer at every pixel of the scan, producing a full mass spectrum for each pixel. The resulting data is a 2D matrix of mass spectra, allowing the extraction of the spatial distribution of ion counts for each detected mass-to-charge ratio (*m/z*) peak (Figs. S1c-d). The mass spectrometer can be operated in positive or negative ion mode, which correspond to voltage settings for extracting either positive or negative secondary ions, respectively. As material is continuously removed by the FIB, scanning the region of interest multiple times enables the recording of three-dimensional (3D) information of the chemical composition of the sample as every 2D image corresponds to a slice orthogonal to the FIB direction, cutting through the sample volume (Fig. S3, Supplementary Video S1). The data collected from cryo-EM and cryo-FIB-SIMS for each region of interest can then be correlatively analyzed to reveal insights into the chemical composition of the sample overlaid with ultrastructural features revealed by cryo-EM (Fig. 1a and Figure S1d).

Initial testing of the FIB-SIMS system on standard chemical samples, such as a vitrified cesium iodide solution (CsI, Fig. S4a) and vitrified water (Fig. S4b) showed all expected ionic peaks. Next, we showed the applicability of the technique to vitrified biological specimens by imaging *Caulobacter crescentus* cells which were coated with gold nanoparticles by labelling of the surface layer (S-layer) protein RsaA^17,18^. The gold nanoparticles are clearly visible in cryo-EM imaging, appearing as black dots around the cell, which indicates successful coating of the entire cell surface (Fig. 1b). Imaging the same cells using cryo-FIB-SIMS reveals a clear gold signal (Au, 197 *m/z*) localized on the cells (Fig. 1b), confirming that our correlative workflow can reveal structural as well as chemical details about the specimen.

### Correlative spatio-chemical imaging of bacterial cells at sub-cellular resolution

To understand the sub-cellular distribution of elements and small molecules within bacterial cells, we imaged unlabelled *C. crescentus* cells (Fig. 2, Figs. S1d, S5) by cryo-EM-FIB-SIMS. The cryo-EM images of the cells clearly show characteristic ultrastructural features, including the inner membrane, outer membrane, S-layer, as well as various intracellular features such as storage granules (Fig. 2). Overlaying the cryo-EM data with FIB-SIMS data corresponding to different *m/z*-peaks and hence to different elemental and small molecular ions shows differential ionic distributions within the cell. Some elements, such as sodium (Na, 23 *m/z*), are spread across the cytosol, whereas others, e.g. magnesium (Mg, 24 *m/z*), are compactly localized within the storage granules. Potassium (K, 39 *m/z*), on the other hand, shows both a strong presence in the cytosol as well as in a clear, prominent peak within the storage granules (Fig. 2a). Interestingly, the smaller granule in Fig. 2a, showing much lighter contrast in cryo-EM, also exhibits a weaker signal in SIMS imaging compared to the larger, darker granule (see arrows in Fig. 2a), indicating a correlation between contrast in cryo-EM and the strength of the signal in SIMS elemental imaging. To supplement these observations about the spatial distribution of various positively charged ions, we subsequently imaged *C. crescentus* cells in negative ion mode. This confirmed that the storage granules are additionally rich in phosphates (PO2 (63 *m/z*), PO3 (79 *m/z*), Fig. 2b), as reported previously^19–22^.

**Figure 2:**
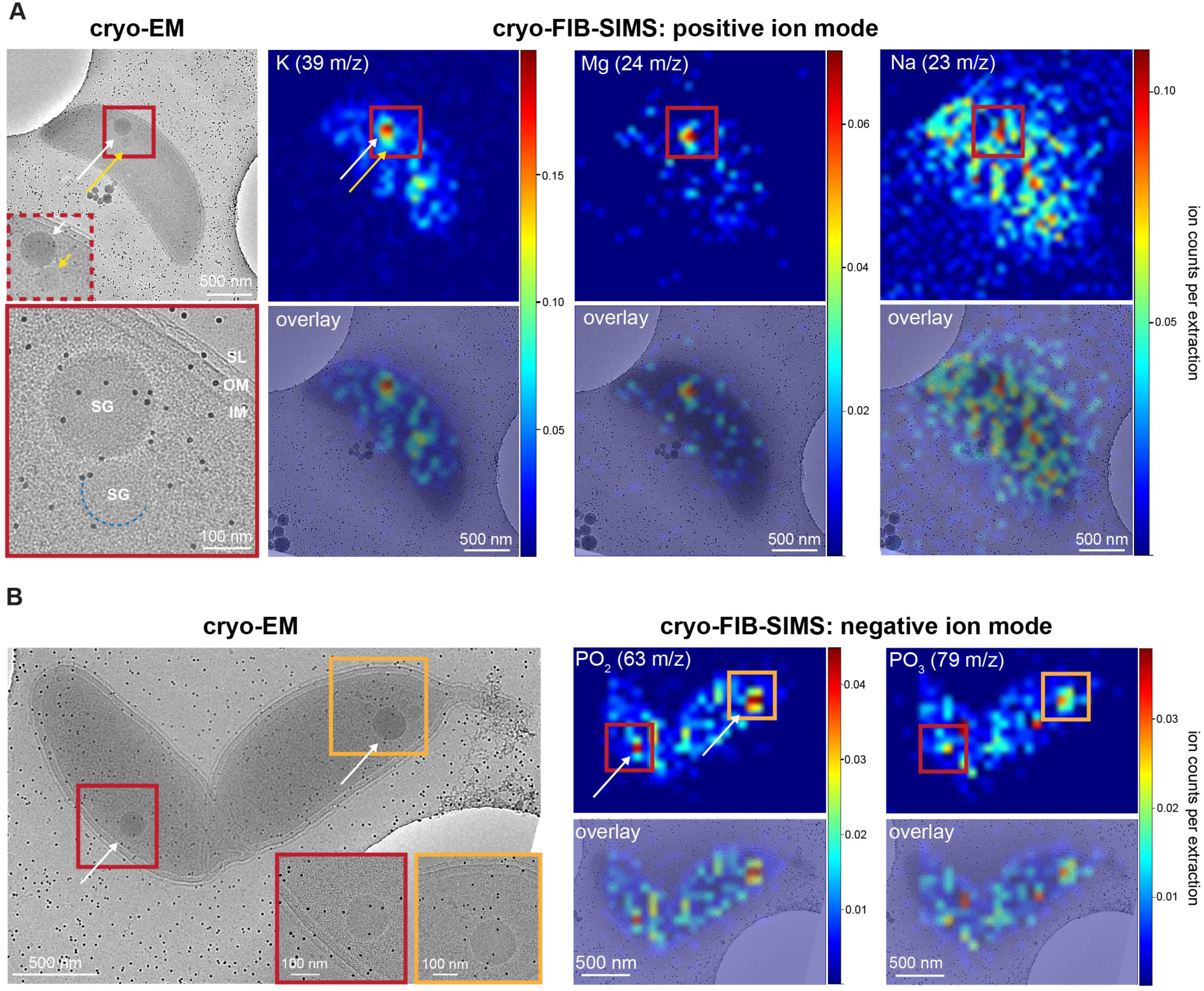
Sub-cellular resolution correlative cryo-EM-FIB-SIMS imaging of bacteria. **(A)** Positive ion mode correlative analysis of a *C. crescentus* cell showing the spatial cryo-EM images at low (top) and high (bottom) magnification as well as cryo-FIB-SIMS images of three prominent *m/z* peaks (K, Mg, and Na) and the respective overlays of spatial and chemical data. The spatial distributions of the different ions in the cell differ significantly. Ultrastructural cellular features such as the inner membrane (IM), outer membrane (OM), S-layer (SL) and storage granules (SG) are labelled in the high-magnification cryo-EM image. For better visualization of the storage granules, which are labelled by a white and yellow arrow, respectively, a zoom-in of the low-magnification image is shown in the inset (red dashed box) and the outline of the smaller granule is enhanced in the high-magnification image (blue dashed line). The black dots are 10 nm protein-A-gold fiducials added to this otherwise untagged specimen. **(B)** Negative ion mode correlative analysis of a *C. crescentus* cell demonstrates the presence of phosphates within the storage granules of the cell.

### Cryo-EM-FIB-SIMS is compatible with cryo-LM as well as multicellular specimen imaging

While cryo-EM-FIB-SIMS supports spatio-chemical imaging of sub-cellular features in bacteria, the field of view of a single measurement is small compared to light microscopy. Integrating cryo-EM-FIB-SIMS into previously established correlative light and electron microscopy (CLEM) workflows^23,24^ would thus allow large field of view imaging for target identification as well as dual labelling strategies featuring both chemical and fluorescent tags. To this end, we extended the experimental system of gold nanoparticle-tagged bacterial cells (Fig. 1b) to include differential fluorescent labelling of the S-layer protein. Using a SpyCatcher-SpyTag system for obtaining fluorescently labelled S-layers^18,25^, we labelled the S-layers either with superfolder GFP or mRFP. In addition, the mRFP-labelled cells were labelled with gold nanoparticles while the GFP-labelled cells were left without gold labelling (Fig. 3a, inset). These differentially labelled cells were mixed prior to vitrification, resulting in an intermingled population of cells that are distinguishable in all three techniques: magenta vs green fluorescence in light microscopy (Fig. 3a), nanoparticle visibility on the cell surface in cryo-EM (Fig. 3b), and detection of gold (Au, 197 *m/z*) signal in FIB-SIMS (Fig. 3c). Imaging the same cells using all three imaging modalities demonstrates the compatibility of the full cryo-CLEM-FIB-SIMS workflow: cells that were mRFP-labelled (magenta) in light microscopy showed extensive gold labelling in both cryo-EM and FIB-SIMS imaging, whereas the green-fluorescent cells did not exhibit gold labelling as confirmed by both cryo-EM and FIB-SIMS (Fig. 3a-d, Fig. S6).

**Figure 3:**
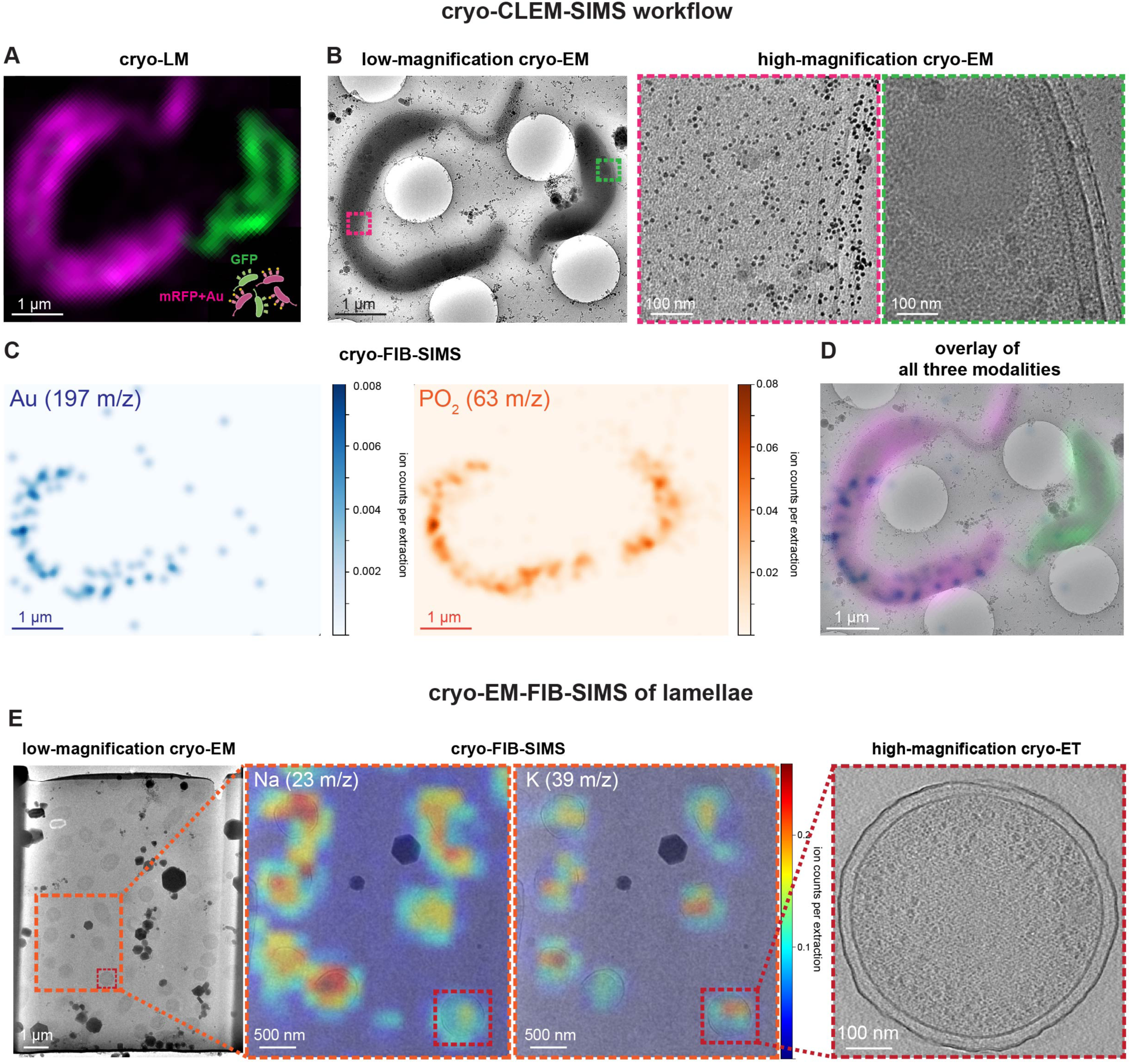
Compatibility of cryo-EM-FIB-SIMS with light microscopy workflows and thick specimen imaging. **(A-D)** Proof-of-principle cryo-CLEM-FIB-SIMS workflow integrating cryo-LM and cryo-EM-FIB-SIMS imaging. **(A)** *C. crescentus* cells with two types of S-layer protein labelling were prepared, resulting in a mixed population of cells with GFP labelling (green) and mRFP plus gold labelling (magenta+Au). **(B)** Imaging the same cells in cryo-EM demonstrates that the magenta-fluorescent cells feature gold nanoparticle labelling (black dots), whereas the green-fluorescent cells do not, as demonstrated both in the low-magnification image as well as in the high magnification images of the regions marked by dashed magenta and green boxes. **(C)** Cryo-FIB-SIMS only detects the magenta+Au cells in the *m/z* channel corresponding to elemental gold (left, blue), while both cells are detected in the PO2 channel (right, orange), which can be further visualized **(D)** by the overlay of all three imaging modalities (cryo-LM image, low-magnification cryo-EM image, cryo-FIB-SIMS image of the Au distribution). **(E)** Cryo-EM-FIB-SIMS workflow applied to cryo-FIB-milled lamellae containing *P. aeruginosa* bacteria and further correlated with cryo-ET of FIB-milled cells.

Many questions of biological interest concern tissues or other multicellular samples, which typically require thinning prior to cryo-EM imaging^26^. Currently, the state-of-the-art approach is cryo-FIB-milling^27,28^, which produces lamellae from a thick specimen via the ablation of material above and below the target of interest. The resulting thin (100-250 nm) lamellae are suitable for imaging by cryo-EM, and specifically by electron cryotomography (cryo-ET)^1,29^, which allows 3D imaging of biological specimens. Given the importance of lamellae for cryo-ET imaging, we sought to apply cryo-FIB-SIMS imaging to lamellae. Using lamellae that were back-coated to improve stability, we imaged a specimen of concentrated *Pseudomonas aeruginosa* bacteria, showing clearly recognizable individual cells in the cryo-FIB-SIMS images (Fig. 3e). These images can be correlated with cryo-EM overview images as well as with high-resolution tomographic slices (Fig. 3e), demonstrating that the cryo-FIB-SIMS workflow is compatible with cryo-ET of lamellae.

### Tracking the accumulation of pollutants in environmental bacteria using cryo-EM-FIB-SIMS

Following the proof-of-principle experiments detailed above, we applied the cryo-EM-FIB-SIMS workflow to the study of pollutant accumulation in environmental bacteria. We studied the uptake of BPAF, a fluorinated environmental contaminant originating from plastics production, which causes endocrinal disruption in humans^30^. Given their fluorinated nature, BPAF molecules can be tracked in our cryo-FIB-SIMS setup due to a strong fluorine signal in negative ion mode (F, 19 *m/z*), which can be used to trace the presence of fluorinated chemicals^16^.

Exposure of the environmental bacterium *C. crescentus* to BPAF lead to a dramatic retardation in growth for several hours (Fig. 4a), along with a significant accumulation of BPAF in exposed cells (Fig. 4b, Fig. S7). Proteomics characterization of cells exposed to BPAF in stationary phase for durations between 15 minutes and 24 hours showed clear overall differences in protein expression relative to untreated cells at all time points (Fig. 4c). BPAF-exposed cells showed retarded proteome changes along the principal component associated with exposure time (Fig. 4c; PC1), especially at 4 and 6 hours. An orthogonal difference in protein expression was observed between BPAF-exposed and unexposed cells (Fig. 4c; PC2). Specifically, we observed upregulation of several drug efflux pumps, including acrB2 and their regulatory proteins, such as TipR (Fig. 4d), indicating that the bacteria attempt to reduce the cytosolic concentration of BPAF by pumping the pollutant out. However, despite the upregulation of drug efflux machinery, bioaccumulation analysis of bacteria exposed to BPAF detected large amounts of BPAF within the bacteria (Fig. 4b).

**Figure 4:**
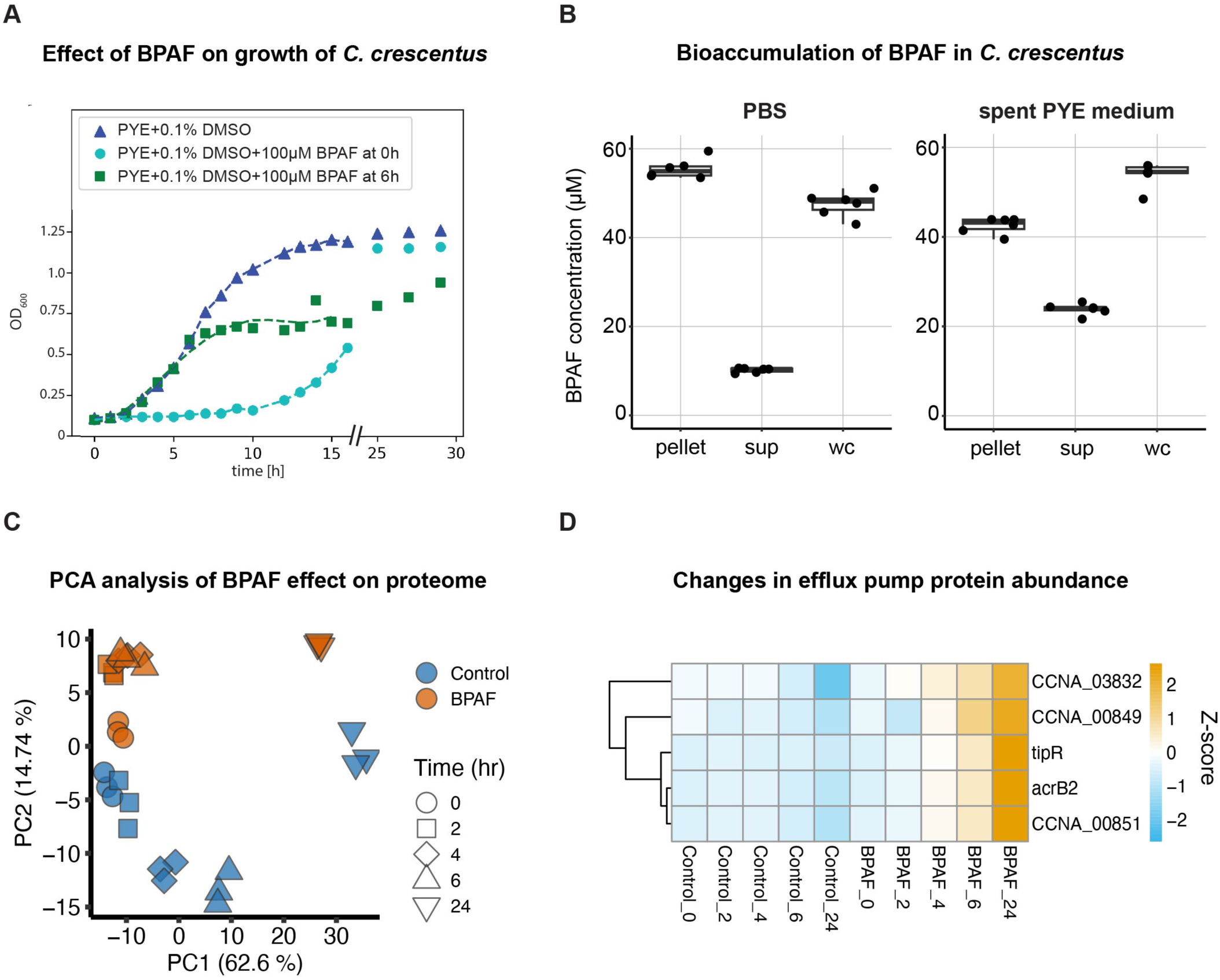
Effect of BPAF on *C. crescentus* cells. **(A)** Growth curves of C. crescentus with (cyan, green) and without (blue) addition of BPAF show that BPAF retards growth for several hours upon addition, both when added at low OD600 at time point 0 hours (cyan) and when added upon reaching exponential growth phase at time point 6 hours (green). **(B)** Bioaccumulation analysis performed both in PBS and spent PYE (Peptone Yeast Extract) medium shows clear differences in BPAF concentration between supernatant and pellet after exposure to 50 µM BPAF, indicating BPAF accumulation within the cells. wc denotes the whole culture (bacteria plus supernatant, PYE or PBS with final concentrations of 50µM BPAF), sup denotes the supernatant (PYE or PBS with 50 µM BPAF after the removal of the bacteria by spinning), and pellet denotes the cell pellet from the wc (exposed to 50 µM BPAF), reconstituted in spent medium or PBS without BPAF. The higher efficiency of bioaccumulation in PBS compared to the spent PYE medium is expected given the lack of nutrients in PBS. Comparison to the internal standard BPS (Fig. S7) demonstrates that the observed pattern is a biological and not a technical effect. **(C)** Proteomics shows that the BPAF exposure leads to large-scale changes in the bacterial proteome as shown by Principal Component Analysis (PCA) of the effect of BPAF treatment for different exposure times. **(D)** Drug efflux machinery and their regulatory proteins are upregulated in BPAF-exposed cells, in a strong bacterial response to keep the intracellular BPAF-concentration low. Efflux pump-associated proteins with a significant change in abundance in BPAF-exposed versus unexposed cells is shown at incubation times 15 minutes (labelled 0), 2 hr, 4 hr, 6 hr, and 24 hr.

While bulk-level bioaccumulation analysis demonstrated uptake of BPAF by *C. crescentus* cells, the resulting intracellular ramifications remained unclear. To address this, we employed our cryo-EM-FIB-SIMS workflow to trace the BPAF-associated fluorine signal in the specimen and correlate it to spatial features observed in cryo-EM. Cryo-EM images of bacteria exposed to 100 µM BPAF for 4 hours showed large cytosolic granular aggregates of different intensities (Fig. 5a, Fig. S8). While the spherical, medium-contrast aggregates (Fig. 5a, red arrows) are likely storage granules frequently observed in *C. crescentus* (Fig. 2, Figs. S1d, 5, and Supplementary Video S2)^19–22^, the high-contrast, often irregularly shaped features (Fig. 5a, blue arrow, Fig. S8, Supplementary Video S2) are not commonly observed in *C. crescentus*. The cryo-FIB-SIMS imaging carried out on the same cells (Fig. 5a-c and Fig. S8) showed a highly localized fluorine signal (Fig. 5b, c) that was not observed in the unexposed control cells (Fig. 5d, e, g), indicating that the localized fluorine signal in the treated cells is due to the presence of BPAF molecules. In contrast to the BPAF signal, other cell-associated signals, such as PO2, were distributed over the entire cell (Fig. 5e, g). The overlay of the cryo-EM and the fluorine cryo-FIB-SIMS images (Fig. 5c) demonstrated that the BPAF accumulation can be traced to a specific type of aggregate, namely to the storage granules, which co-localize with the fluorine signal. In contrast, the cytosol as well as the other types of aggregates do not exhibit localized fluorine signal (n=162 cells, from two biological replicates).

**Figure 5:**
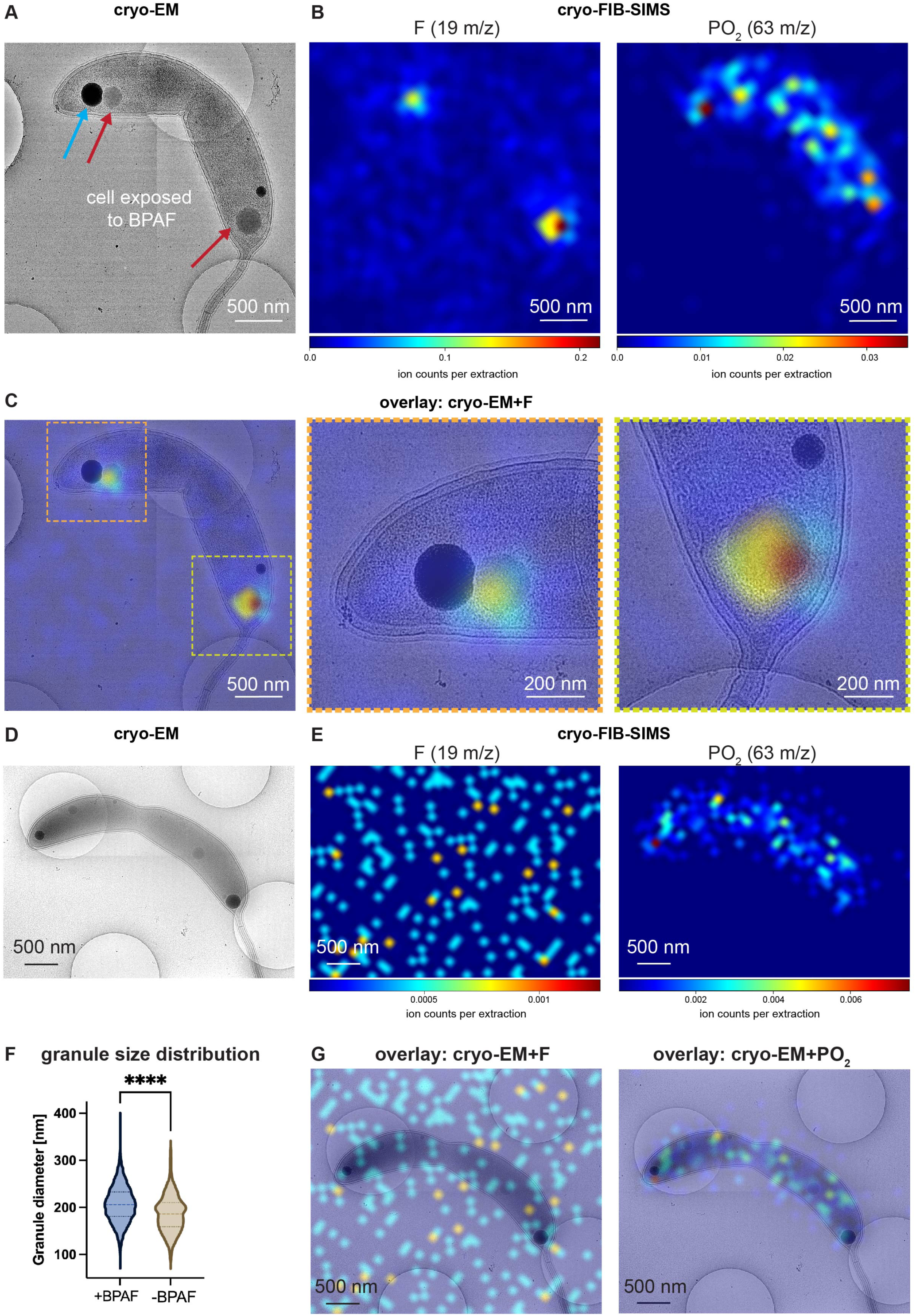
Intracellular localization of BPAF in *C. crescentus* cells. **(A)** Cryo-EM image of a *C. crescentus* cell after 4 hours exposure to 100 μM BPAF. Two different types of aggregates are observed within the cell: high-contrast (black) aggregates (blue arrow) and medium-contrast spherical aggregates reminiscent of storage granules (red arrows, compare with Fig. 2, Figs. S1, S5). **(B)** Cryo-FIB-SIMS images of the same cell as in **(A)** for the *m/z* peaks 19, corresponding to fluorine (left), and 63, corresponding to PO2 (right). While the PO2 signal is detectable over the entire cell, the fluorine signal, marking the presence of BPAF, is highly localized. **(C)** Overlay of the fluorine signal and the cryo-EM image at medium magnification (left) and high magnification (right) shows that the fluorine signal is localized in the storage granules, showing that BPAF is accumulated in these aggregates. **(D)** Cryo-EM image of a *C. crescentus* cell from the same culture as the cells presented in Fig. 5 and Fig. S8, grown for 4 hours in PYE media with 0.1% DMSO without BPAF (unexposed control sample). **(E)** Corresponding cryo-FIB-SIMS signals of fluorine (F, 19 *m/z,* lef*t*) and PO2 (63 *m/z,* right). The fluorine signal is several orders of magnitude lower than the signal observed in BPAF-exposed-cells and does not show any specific localization in the cell. **(F)** Comparison of the storage granule diameter in cells exposed to BPAF (left, blue) and unexposed control cells (right, beige) shows a statistically significant (p < 0.0001) increase in granule size. **(G)** Overlay of cryo-EM image and the cryo-FIB-SIMS images of fluorine (left) and PO2 (right).

While both the BPAF-exposed and the unexposed cells (Fig. 5f) show a large range of storage granule sizes, the mean diameter of the granules in the exposed cells shows a statistically significant (p < 0.0001) increase (206.9 ± 38.5 nm, n=1,464) compared to the unexposed control cells (186.4 ± 37.9 nm, n=252). These results strongly suggest that BPAF is sequestered in the granules after uptake by the cells, thus removing BPAF from the cytosol. In conjunction with the upregulation of drug efflux machinery, the sequestration of BPAF in the granules suggests that BPAF cannot be efficiently removed from the cell, in line with observations in *E. coli* Δ*tolC* mutants described previously^16^ for other fluorinated compounds. This indicates that sequestration inside aggregates could be an alternative way of lowering pollutant concentration in the cytosol. Imaging BPAF-exposed cells at both shorter (2 hours) and longer exposure times (24 hours) showed strongly localized intracellular aggregation of BPAF at both time points (Fig. S9), implying that the pollutant is taken up quickly and retained within the cells for a long period of time. Cryo-EM-FIB-SIMS imaging thus allowed us to pinpoint the precise localization of BPAF accumulation within *C. crescentus* cells, yielding insights into how these cells accumulate toxic environmental pollutants at the sub-cellular level.

## Discussion

We report a cryo-EM-FIB-SIMS workflow (Figs. 1-2) that can be applied in a modular way to a variety of samples, including untagged cells (Fig. 2), fluorescently tagged cells (Fig. 3a-d) and multicellular specimens (Fig. 3e). The workflow is also compatible with cryo-ET and could hence be developed further for correlative cryo-ET-FIB-SIMS imaging, as well as for 3D cryo-FIB-SEM (serial block face imaging), i.e. volumetric imaging.

While the use of a monoatomic Ga beam allows SIMS imaging at relatively high spatial resolution^4,31^, thus enabling the visualization of ∼100 nm storage granules (Fig. 2a), the low ionization yield, as well as the large interaction depth of the Ga beam with the sample^32^, results in a very low probability of extracting peptides, lipids or oligonucleotides. To detect these larger ions, specific metal-labelling^12^ or isotope-labelling of specimens^33–35^ could be used. Another possibility to detect larger molecules would be to utilize larger ion beams^9,11,12^, including gas cluster ion beams (GCIB)^6^.

We used our workflow to study the outstanding question of the uptake of fluorinated contaminants by environmental bacteria, demonstrating the power of correlative spatio-chemical imaging for uncovering mechanistic biology. We suggest that such combinations of techniques with high spatial and chemical resolution, such as cryo-EM-FIB-SIMS described here, along with other recently reported methods^15,36–39^ will be needed to understand other complex biological specimens at the molecular mechanistic level.

## Acknowledgements

This work was supported by the Medical Research Council, as part of United Kingdom Research and Innovation (also known as UK Research and Innovation) [Programme MC_UP_1201/31 to T.A.M.B]. For the purpose of open access, the MRC Laboratory of Molecular Biology has applied a CC BY public copyright license to any Author Accepted Manuscript version arising. T.A.M.B. would like to thank the Wellcome Trust (grants 225317/Z/22/Z and 223788/Z/21/Z) and the Lister Institute for Preventative Medicine for support. H.O. was supported by an EMBO Long-term fellowship (ALTF 1076-2023). The authors would like to thank Stephan Rauschenbach and Christopher J. Russo for helpful discussions and Andrew Carter, Sjors Scheres, Ido Caspy and Abul Tarafder for critical reading of the manuscript. The authors also gratefully acknowledge the MRC LMB electron microscopy, light microscopy and mass spectrometry facilities.

This paper was typeset with the bioRxiv word template by @Chrelli: www.github.com/chrelli/bioRxiv-word-template

## Author contributions

This study was conceived by T.A.M.B., H.O. and K.R.P. Experiments and data analysis were performed by H.O., B.I., S.B., D.S., Z.W., Y.Z., C.F., R.B. and T.S. Funding was acquired by T.A.M.B., K.R.P. and H.O. T.A.M.B. and K.R.P. supervised the project. The paper was drafted by H.O, B.I., and T.A.M.B. All authors contributed to the discussion and finalization of the paper.

## Data availability

All data are either available in the main text and the Materials and Methods or available upon request. The mass spectrometry proteomics data have been deposited to the ProteomeXchange Consortium via the PRIDE partner repository.

## Materials and Methods

### Preparation of CsI and H2O samples for cryo-FIB-SIMS

Vitrified samples of CsI and H_2_O were prepared by applying 2.5 µl of saturated CsI solution (in water) and deionized water, respectively, to cryo-EM grids (Quantifoil Cu/Rh R3.5/1), followed by blotting from both sides in a Vitrobot Mark IV (ThermoFisher) and plunge-freezing in liquid ethane which was maintained below -170 °C. The chamber was kept at 100% relative humidity at a temperature of 10 °C, with blotting conditions comprising a blot time of 5 s, blot force of -10, and wait time of 2 s.

### Unlabeled *C. crescentus* for cryo-EM-FIB-SIMS analysis

*C. crescentus* cells (strain CB15N) were inoculated from frozen stock into 5 ml of PYE (Peptone Yeast Extract) medium (0.2% (w/v) bactopeptone, 0.1% (w/v) yeast extract, 0.5 mM CaCl_2_, 1 mM MgSO_4_,^40^) and grown in a shaking culture at 30 °C to an optical density measured at 600 nm wavelength of light (OD_600_) of 1. Subsequently, cells were prepared for cryo-EM imaging as described previously^41^. Briefly, 2.5 µl of cell culture was applied to a cryo-EM grid (Quantifoil Cu/Rh R1.2/1.3), blotted from both sides in a Vitrobot Mark IV (ThermoFisher) and plunge-frozen into liquid ethane which was maintained below - 170 °C. The chamber was kept at 100% relative humidity at a temperature of 10 °C, with blotting conditions comprising a blot time of 2.5 s, blot force of -7, and wait time of 30 s.

### Fluorescent and gold-nanoparticle labelled *C. crescentus* for cryo-CLEM-FIB-SIMS analysis

Purification of SpyCatcher fusion proteins: His-tagged SpyCatcher fusion proteins were purified as previously described^42,43^. In short, plasmids pDEST14-SpyCatcher-superfolderGFP (sfGFP) and pBAD-SpyCatcher-mRFP1 were transformed (separately) into *E. coli* BL21 DE3 cells and grown in LB in the presence of 100 μg/ml ampicillin. For each construct, a 6 L culture was grown at 37 °C with shaking until an OD_600_ of 0.8 was reached. The culture expressing SpyCatcher-sfGFP was induced with 0.5 mM IPTG (isopropyl β-D-1-thiogalactopyranoside); while the culture expressing SpyCatcher-mRFP1 was induced with 0.1 % (w/v) arabinose for 16 hours at 18 °C. Cells were pelleted at 4,000 rcf (relative centrifugal force) for 30 minutes at 4 °C and resuspended in 150 ml lysis buffer (50 mM HEPES(4-(2-hydroxyethyl)-1-piperazineethanesulfonic acid)/NaOH pH=7.5, 500 mM NaCl, 1 mM MgCl_2_, 5% Glycerol, 50 µg/mL DNaseI, 50 µg/mL RNaseA, 0.2 mM TCEP (tris(2-carboxyethyl)phosphine), 2 complete Protease Inhibitor tablets (Roche)). Cells were passed through a homogenizer at a pressure of 20,000 psi (pounds per square inch) for two cycles. Cell debris was pelleted at 20,000 rcf for 30 minutes at 4 °C. The supernatant was clarified by passing through a 0.22 μm filter and loaded onto a 5 ml HisTrap column (Cytiva). Elution was performed by applying the elution buffer (50 mM HEPES/NaOH pH=7.5, 500 mM NaCl, 5% glycerol, 0.2 mM TCEP, 500 mM imidazole) with a shallow gradient. The fractions containing the SpyCatcher conjugates were pooled and dialysed overnight into the gel filtration buffer (50 mM HEPES/NaOH pH=7.5, 150 mM NaCl, 5% glycerol, 0.2 mM TCEP). The protein solution was concentrated to a 5 ml volume and then loaded onto a HiLoad Superdex S200 16/600 column (Cytiva). The eluted peaks were pooled, flash-frozen in liquid nitrogen and stored at -80 °C until further use.

*C. crescentus* labelling with SpyCatcher fusion proteins and NanoGold: *C. crescentus* cells (CB15N *ΔsapA rsaA467*:SpyTag) were grown in PYE (Peptone Yeast Extract) at 30 °C with shaking at 180 rpm (revolutions per minute). Cells grown to exponential growth phase were diluted to an OD_600_ of 0.1 in a total culture volume of 250 μl and mRFP-SpyCatcher was added to final concentration of 8 μM. The labelling mixture was incubated at 23 °C for 90 minutes. After the incubation, cells were washed three times with PYE and pelleted at 6,000 rcf for 3 minutes at 23 °C. Cells were resuspended into 105 μl PYE after the final wash, to which 105 μl of 5 nm-Ni-NTA (nickel (II) nitraloacetic acid) NanoGold (Nanoprobes) was added to achieve a final 0.25 μM concentration of 5-nm-Ni-NTA-NanoGold. The final NaCl concentration was brought to a more physiologically relevant range of 120 mM. After this, the gold labelling reaction was commenced by incubation for 30 minutes at 23 °C. After incubation, cells were washed three times with PYE and resuspended in fresh 5 μl PYE. In parallel, cells labelled with sfGFP-SpyCatcher were prepared using the same procedure except for the omission of the gold labeling reaction. Briefly, cells grown to exponential phase were diluted to an OD_600_ of 0.1 with a final volume of 250 μl and sfGFP-SpyCatcher was added to a final concentration of 8 μM and incubated for 90 minutes at 23°C. Cells were washed three times, and the final resuspension was performed into 5 μl of PYE. Prior to plunge-freezing for cryo-EM, mRFP-SpyCatcher-Nanogold and sfGFP-Spy-Catcher labelled cells were mixed in a 2:1 ratio.

Cryo-EM grids were prepared as described previously^44^. Briefly, 3.5 µl of the resuspended sample was applied to a freshly glow discharged 135 mesh Quantifoil London Finder grid (H15) and plunge-frozen into liquid ethane maintained at -178 °C, using a Vitrobot Mark IV at 10 °C and relative humidity of 95% after a wait time of 30 s, with -2 blot force, 0.5 s drain time and 2.5 s blot time.

### High-pressure frozen *P. aeruginosa* for cryo-EM-FIB-SIMS of lamellae

*P. aeruginosa* (strain PAO1) cells were cultured overnight on an LB agar plate. Bacterial cells were harvested by scraping off the plate and resuspended in LB media with 5% glycerol to reach an OD_600_ of around 100. Cells were vitrified on an EM grid by high-pressure freezing using a Compact 03 HPF (M. Wohlwend GmbH) following the “waffle” method protocol^45^.

### Sample preparation for studying BPAF uptake by *C. crescentus* using cryo-EM-FIB-SIMS and proteomics

*C. crescentus* cells were inoculated from a frozen glycerol stock culture into 30 ml PYE medium and grown overnight at 30 °C. The resulting pre-culture was then used to inoculate three flasks with 400 ml PYE medium (10 ml per flask) and grown for 24 hours. The culture was then centrifuged at 4,500 rcf for 10 minutes to pellet the cells. The cells were next resuspended into the same medium and adjusted to an OD_600_ of 3.75 for cryo-EM-FIB-SIMS imaging and to an OD_600_ of 2 for proteomics studies. For cryo-EM-FIB-SIMS, the culture was split into two flasks containing 30 ml culture each. To one of the two flasks, a stock solution of BPAF in DMSO (dimethyl sulfoxide) was added resulting in a final concentration of 100 µM BPAF and 0.1% (v/v) DMSO. To the other flask, serving as a control sample, 0.1% (v/v) DMSO was added. Both flasks were incubated at 30 °C for 4 hours and two independent biological replicates were analyzed. The cells were prepared for cryo-EM analysis as described in the previous section for unlabeled cells. For proteomics, the culture was split into 6 flasks of 30 ml culture each. To three of them, BPAF in DMSO was added to achieve a final concentration of 100 µM BPAF and 0.1% (v/v) of DMSO, and the control samples were adjusted to a final concentration of 0.1% (v/v) DMSO. Samples were collected after 15 mins and after 2, 4, 6, and 24 hours in a standing incubation at 30 °C. Three ml of each of the samples were centrifuged at 16,800 rcf for 1 minute and the pellet was directly frozen at -80 °C.

### Bioaccumulation analysis of BPAF in *C. crescentus*

*C. crescentus* growth and treatment: *C. crescentus* cells were grown overnight in PYE medium, the culture was split, centrifuged at 4,500 rcf for 10 minutes, and the pellets were washed once with PBS and spent PYE media (the media in which the cells were grown in), respectively, and subsequently resuspended at an OD_600_ of 3.75 in the respective media (PBS or spent PYE). 50 µM BPAF were added to the samples from a 50 mM stock solution in DMSO. The samples (6 replicates, three DMSO controls and four incubation controls without bacteria) were incubated at 30 °C for 4 hours.

Sample collection and extraction for BPAF bioaccumulation: Whole culture (wc) was harvested and the remaining sample was centrifuged for 2.5 minutes at 30 °C at 16,900 rcf. Supernatants (sup) were collected and the pellets were resuspended in equivalent amount of BPAF-free PBS and spent PYE media, respectively. All samples were extracted by adding 4x the amount of ice cold (-20 °C) extraction buffer (1:1 ratio of methanol and acetonitrile with internal standards, 50 µM bisphenol S (for negative mode), spiked in) followed by an incubation on ice for 15 min. Samples were then centrifuged at 4 °C at 16,900 rcf for 2.5 minutes and supernatants were transferred to vials for liquid-chromatography-mass-spectrometry (LC-MS) analysis. Samples for concentration calibration (1 in 2 dilution of BPAF) and bacteria-free compound controls were processed in the same way.

LC-MS measurements for BPAF bioaccumulation: Analysis was performed on an Agilent 1290 Infinity II LC system coupled with an Agilent 6470 triple quadrupole mass spectrometer with JetStream ESI source operated in dynamic multiple reaction monitoring (dMRM) mode. BPAF and bisphenol S were detected in negative ion mode. Chromatographic separation was performed using a ZORBAX RRHD Eclipse Plus column (C18, 2.1 × 100 mm, 1.8 µm; Agilent 858700-902) at 40 °C. The multisampler was maintained at a temperature of 4 °C. The injection volume was 1 µl and the flow rate was 0.4 ml/minute. The mobile phases consisted of A: water + 0.1 % formic acid + ammonium formate 10 mM; B: acetonitrile + 0.1 % formic acid + ammonium formate 10 mM. The 10 min gradient started with 35 % solvent B, which was increased to 100 % by 9 minutes and held for one minute before returning to 35 %. A pooled QC sample, blanks and serial dilutions of BPAF were injected regularly throughout each run. Data was analysed using MassHunter Workstation Quantitative Analysis for QQQ v10.1. Concentrations were obtained via external calibration using standard curves.

### Cryo-FIB-SIMS

Cryo-FIB-SIMS measurements were carried out on a gallium FIB-SEM (Zeiss Crossbeam 550) equipped with a ToF mass spectrometer (TOFWERK fibTOF). During cryo-FIB-SIMS imaging, the region of interest was scanned by the gallium FIB, whose interaction with the sample results in the ablation of material at each pixel. The resulting secondary ions were extracted into and analyzed by the ToF mass spectrometer at each pixel. As a full mass spectrum is produced at each pixel, the chemical composition of the sample can be visualized by plotting the pixel-by-pixel intensity for each detected mass-to-charge ratio. The range of detectable mass-to-charge ratios depends on the dwell time of the FIB at each pixel. For the standard settings used here (FIB dwell time per pixel 12.8 μs), this corresponds to a *m/z*-range from 1 to 323. The image intensity (colour scale) maps the ion counts per extraction (one extraction corresponding to one pixel in FIB scanning, see Fig. S1). Simultaneously, secondary electrons resulting from the interaction of the FIB and the specimen can be detected, yielding spatial images of the sample (FIB images, see Fig. S2). Depending on the polarity of the extraction voltage, either positively or negatively charged secondary ions can be extracted and detected. As the imaging process removes material from the sample, repeated scanning of the area of interest yields 3D information of the sample composition as each 2D data set can be regarded as a slice of the 3D sample volume analogous to a ‘Z’-stack in light microscopy, where the ‘Z’ axis is the ion-beam direction. The resulting volume hence allows both extraction of ‘X-Y’-view images (frames) orthogonal to the ion beam direction as well ‘X-Z’ or ‘Y-Z’ slices along the direction of the ion beam for visualizing the variation of sample composition with depth (Fig. S3 and Supplementary Video S1).

For cryo-FIB-SIMS imaging, the Ga beam was operated at 30 kV and 50 pA, the scanning area was 512 pixel by 512 pixel with mass spectrometry data collected using a binning of 8 pixel x 8 pixel, a FIB dwell time per pixel of 12.8 µs, extraction pulse with of 1000 ns and magnification-dependent unbinned pixel sizes ranging from 10-30 nm. The depth of material removed by a single scan (corresponding to one frame) depends on beam current, pixel size and dwell time. For the beam settings reported here, a rough estimate using the sputtering rate of Ga beam milling on vitrified ice^46^ yields a depth per frame of 10-50 nm. The mass resolution of the instrument is specified as >700 by the manufacturer and was measured to be in the range of 1000-1500 in the experiments performed during this study.

The following numbers of cells were analyzed in the cryo-FIB-SIMS experiments shown in this study: Gold nanoparticle labelled *C. crescentus* (Fig. 1b) n=24; unlabelled *C. crescentus* (Fig. 2): n=105; cryo-CLEM-FIB-SIMS: mRFP+Au-labelled (Fig. 3): n=24, GFP-labelled: n=29; BPAF uptake (Fig. 5): n=162 BPAF-exposed, n=50 unexposed. Storage granule size measurements have been carried out in Fiji ^47^ from cryo-EM images for n=1,464 granules in BPAF-exposed cells, n=252 in unexposed cells, and statistical tests (unpaired t-test) have been performed using GraphPad Prism version 10.4.1.

For the cryo-FIB-SIMS experiments on lamellae, 15 lamellae of concentrated *P. aeruginosa* were imaged.

### Sample backside coating for enhanced stability-FIB-SIMS analysis

For compatibility with cryo-EM imaging, specimens need to be prepared on electron-transparent supports such as grids with amorphous carbon films. In principle, cryo-FIB-SIMS imaging can be directly carried out on these specimen supports^16^, however, for increased stability and reduced sample movement, which increases both the overall MS imaging quality and specifically the signal intensity from holey supports (see Fig. S2), we found it beneficial to apply an organometallic coating prior to cryo-FIB-SIMS imaging. For this, we first sputtered the back of the sample (facing away from the ion beam during cryo-FIBSIMS imaging) with a thin layer of platinum in the transfer load lock (Quorum) and then applied an organometallic coating (C_9_H_16_Pt) to the back of the sample using the Gas Injection System (GIS) in the FIB-SEM chamber (Zeiss).

### Cryo-EM data collection

2D cryo-EM data was collected on either a Titan Krios G1 or a Glacios microscope (ThermoFisher) operating at an acceleration voltage of 300 kV or 200 kV respectively, equipped with a direct electron detector (Gatan/ThermoFisher). Medium-magnification (full cell) images were recorded at nominal magnifications ranging from 6500x to 11500x, corresponding to pixel sizes of 23.89 to 18.01 Å, at exposure times of 2-5 s. High-magnification images were recorded at 42000x magnification (pixel size of 1.856 Å). Cryo-ET data was collected on a Titan Krios G3 microscope (ThermoFisher) operating at an acceleration voltage of 300 kV, fitted with a Quantum energy filter (slit width 20 eV) and a K3 direct electron detector (Gatan). CryoET tilt series were collected using a grouped dose-symmetric tilt scheme as implemented in SerialEM^48^, with a total dose of 120 electrons/Å^2^ per tilt series, nominal defocus of -8 µm, and with ±60° tilts of the specimen stage at 1° tilt increments. Tilt series images were collected using a physical pixel size 3.42 Å.

### Cryo-LM

Fluorescence microscopy images of the vitrified cells were acquired using a ZEISS LSM 900 confocal microscope equipped with an Airyscan 2 detector, Axiocam 306 camera, a Colibri 5 LED light source (ZEISS Microscopy) and a CMS196V3 (serial number DV5260-0009) cryo-stage (Linkam Scientific). Vitrified grids were initially inspected using a 0.5x/0.2 Numerical Aperture (NA) objective and high-resolution images were obtained using an LD EC Epiplan-Neofluar 100x/0.75 NA objective. Due to variable ice thickness, cells were positioned at different depths within the specimen and therefore ‘Z’-stacks of 2 μm thickness with 0.5 μm intervals were acquired. Airyscan processing was performed for all high-resolution images in the Zen software (ZEISS Microscopy). When necessary, image registration was applied using Linear Stack Alignment with the SIFT plugin^49^. Maximum intensity projections were performed on the Z-stacks using Fiji^47^.

### Lamella preparation, cryo-ET data acquisition and visualization

Lamellae of *P. aeruginosa* cells were prepared using a Crossbeam 550 (Carl Zeiss Ltd) equipped with a Quorum cryo-stage following the “waffle” milling strategies reported previously ^45^. Low-magnification images (at 2,300x (pixel size 39.23 Å) or 8700x (pixel size 10.26 Å) nominal magnification), as well as tomographic data were acquired in a Titan Krios G3 (Thermo Fisher Scientific) equipped with a K3 summit energy filter (Gatan). Tilt series were acquired at 42000x nominal magnification (pixel size 2.13 Å) with a dose-symmetric tilt scheme (3° increment, -54° to +54° tilt range relative to the lamella plane). Tomograms were reconstructed using the AreTomo^50^ software and visualized using the IMOD software^51^.

### Proteomics

For mass spectrometry analysis, total protein extracts were prepared by lysing cells in 20 mM HEPES containing 0.2% RapiGest (Waters, w/v). Nucleic acids were digested by incubating the cell lysates with 500 units of benzonase nuclease (Sigma) for 30 min at room temperature. Cysteines were reduced by adding DTT to a final concentration of 4 mM and then alkylated by adding iodoacetamide to a final concentration of 14 mM. Two hundred micrograms of total protein per sample were digested for 16h at 37C using a 50:1 protein:trypsin ratio. For the TMT internal bridge channel, a pooled control containing equal protein amount from all samples was prepared. One-hundred micrograms of protein per sample and the internal control pooled sample were labeled with TMTPro reagents according to manufacturer instructions in a total of two TMTPro plex pooled samples. Multiplexed samples were dried to completion, desalted and basic reverse phase was used to fractionate samples into 12 final concatenated fractions per plex. Samples were analyzed using a Thermo Vanquish Neo (Thermo Scientific, Hemel Hempstead) hyphenated to a Thermo Orbitrap Ascend (Thermo Scientific). Approximately 1 μg of protein per fraction was loaded on a trapping column (Thermo Scientific, PepMap100, C18, 300 µm x 5 mm) and resolved on the analytical column (Aurora Ultimate XT 25cm C18 from IonOpticks) at a flow rate of 300 nl/min using a gradient of 97% A (0.1% formic acid) 8% B (80% acetonitrile 0.1% formic acid) to 28% B over 90 minutes, then to 40% B for additional 15 minutes. Data was acquired using three FAIMS (High-field asymmetric-waveform ion mobility spectrometry) cv’s (-40v, - 50v, -60v) and each FAIMS experiment had a maximum cycle time of 1.2, 1, 1s respectively. Data-dependent synchronous precursor selection with real-time MS3 trigger (SPS-MS3 RTS) was used for data acquisition and consisted of a 120,000 resolution full-scan MS scan (AGC set to 100% (4e5 ions) with a maximum fill time of 50 ms) using a mass range of 400-1400 *m/z*. To avoid repeated selection of peptides for MS/MS, the program used a 60 s dynamic exclusion window. MS/MS was performed on the ion trap at normal scan rate [AGC set to 200% (2xe^4^ ions) with a maximum fill time of 35ms] with an isolation window of 0.7 *m/z* and an HCD collision energy of 33%. Real time search parameters were set as follows: *C. crescentus* Uniprot database (UP000001364, downloaded in 02.25) with trypsin set as the main enzyme, static modifications of cysteine carbamidomethylation (deltaMass=57.0215) and TMTpro 16plex (deltaMass=304.2071) on lysines. Methionine oxidation was set as variable modification (deltaMass 15.99491). One missed cleavage was allowed and FDR filtering was enabled. Only 5 peptides per protein were allowed per basic reverse phase fraction (using the close-out option) and RTS search maximum search time was set to 35ms. Ten of the most abundant peptide fragments were selected for SPS MS3 and MS3 acquired at 120,000 resolution on the mass range 100-500 *m/z* with an AGC target of 200% and a maximum injection time of 2451 ms. MS3 collision energy was set to 55%. Raw data were imported and data processed in Proteome Discoverer v3.1 (Thermo Fisher Scientific). The raw files were submitted to a database search using Proteome Discoverer with SequestHF against the *C. crescentus* Uniprot database (UP000001364, downloaded in 02.25). Common contaminant proteins (several types of human keratins, BSA and porcine trypsin) were added to the database. The spectra identification was performed with the following parameters: MS accuracy, 10 p.p.m.; MS/MS accuracy of 0.6 Da for spectra acquired in Ion trap mass analyzer; up to two missed cleavage sites allowed; carbamidomethylation of cysteine and TMTpro 16 plex on lysines and peptide N-terminal as a fixed modification; and oxidation of methionine as variable modifications. Percolator node was used for false discovery rate estimation and only rank 1 peptide identifications of high confidence (FDR<1%) were accepted.

Peptide-Spectrum match (PSM) level output from Proteome Discoverer was processed in R (v 4.4.1) using QFeatures (v 1.14.2)^52^ and biomasslmb (v 0.0.1) R packages^53^. The R markdown notebooks are available from https://github.com/lmb-mass-spec-compbio/BPAF_uptake_Caulobacter_crescentus_TMT_proteomics and archived by zenodo. PSMs were filtered to remove matches to contaminants and median normalized. PSMs with signal-to-noise ratios < 10, co-isolation > 50%, peptide matching rank < 1 or containing any missing values were removed. Finally, PSMs for proteins with fewer than 2 PSMs were removed, before summarization to protein level abundances by summing PSM-level intensities. The protein abundances from the two TMT plexes were then combined and normalized using the internal reference standard samples^54^, with proteins present in just one plex removed, leaving 2986 proteins quantified across all samples. Statistical testing to compare BPAF-exposure to control samples at each timepoint was perfomed with limma (v3.60.6)^55^ using the treat function to set the null hypothesis as fold-change < 1.25 and allowing a trend between mean protein abundance and the prior variance. P-values were adjusted using the Benjamini-Hochberg FDR procedure^56^ and a threshold of 0.01 (1% FDR) used to identity statistically significant differences. PCA was performed using the *prcomp* function in R, with *centre=TRUE*. Heatmap visualization of efflux-pump proteins was performed using pheatmap (v1.0.12) with proteins Z-score normalized across the rows.

---

**Figure S1:**
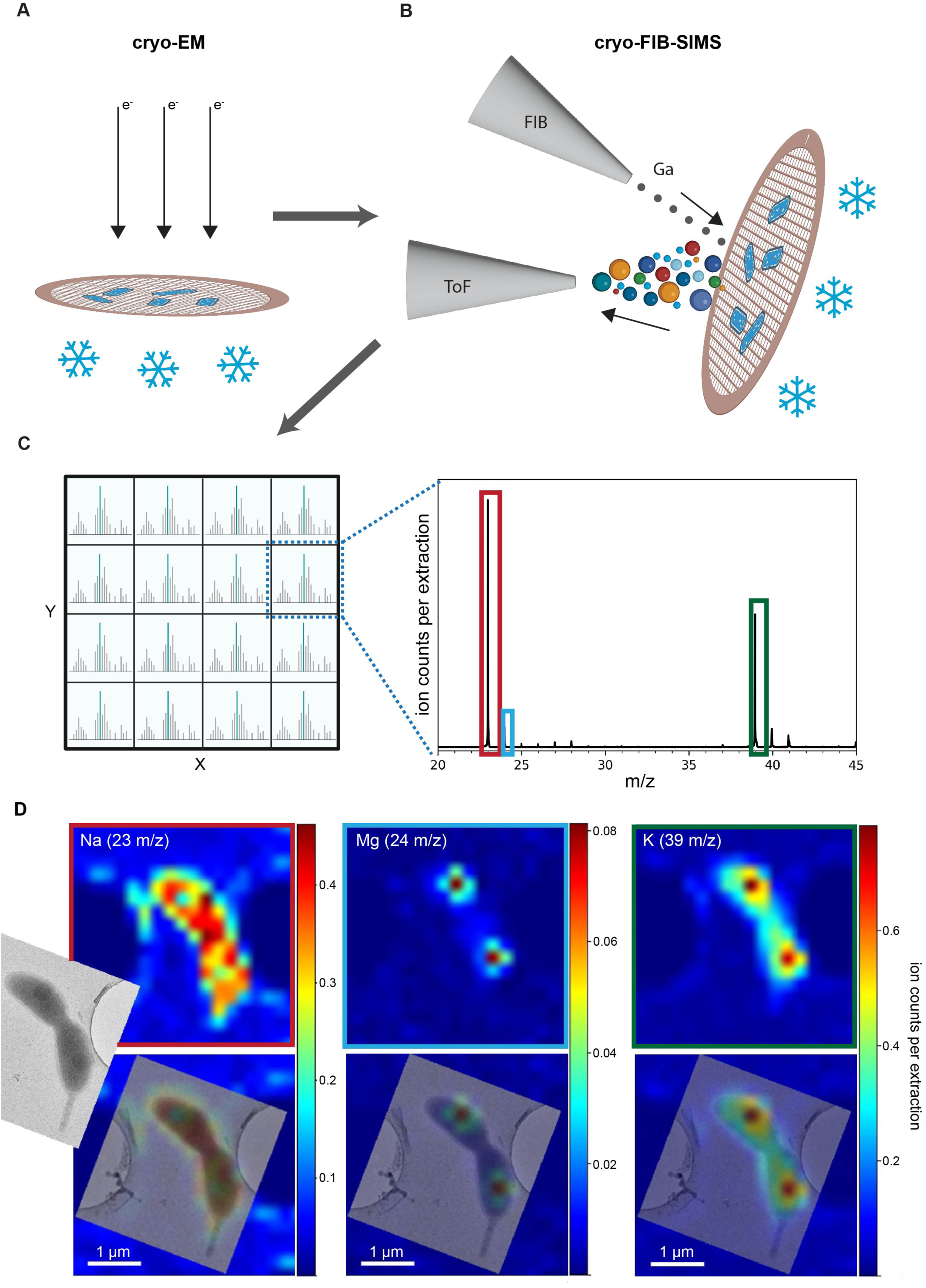
Cryo-EM-FIB-SIMS workflow. **(A-D)** Cryo-EM-FIB-SIMS working principle: after cryo-EM imaging **(A)**, the same specimen area is imaged by cryo-FIB-SIMS. The gallium FIB scans the sample, ablating material during specimen interaction **(B)**. At each pixel, the resulting secondary ions are extracted into the ToF mass spectrometer, yielding a mass spectrum at every pixel **(C)**. From this, a 2D matrix of mass spectra, i.e. the spatial distribution of each detected mass-to-charge ratio (*m/z*) peak can be extracted, as exemplified for a *C. crescentus* cell in **(D)**. Positive ion mode correlative analysis of a *C. crescentus* cell showing the cryo-FIB-SIMS data of three prominent *m/z* peaks (Na, Mg, and K, top) as well as overlays with the cryo-EM image (bottom). The colour scale of the MS images corresponds to the detected ion count per extraction.

**Figure S2:**
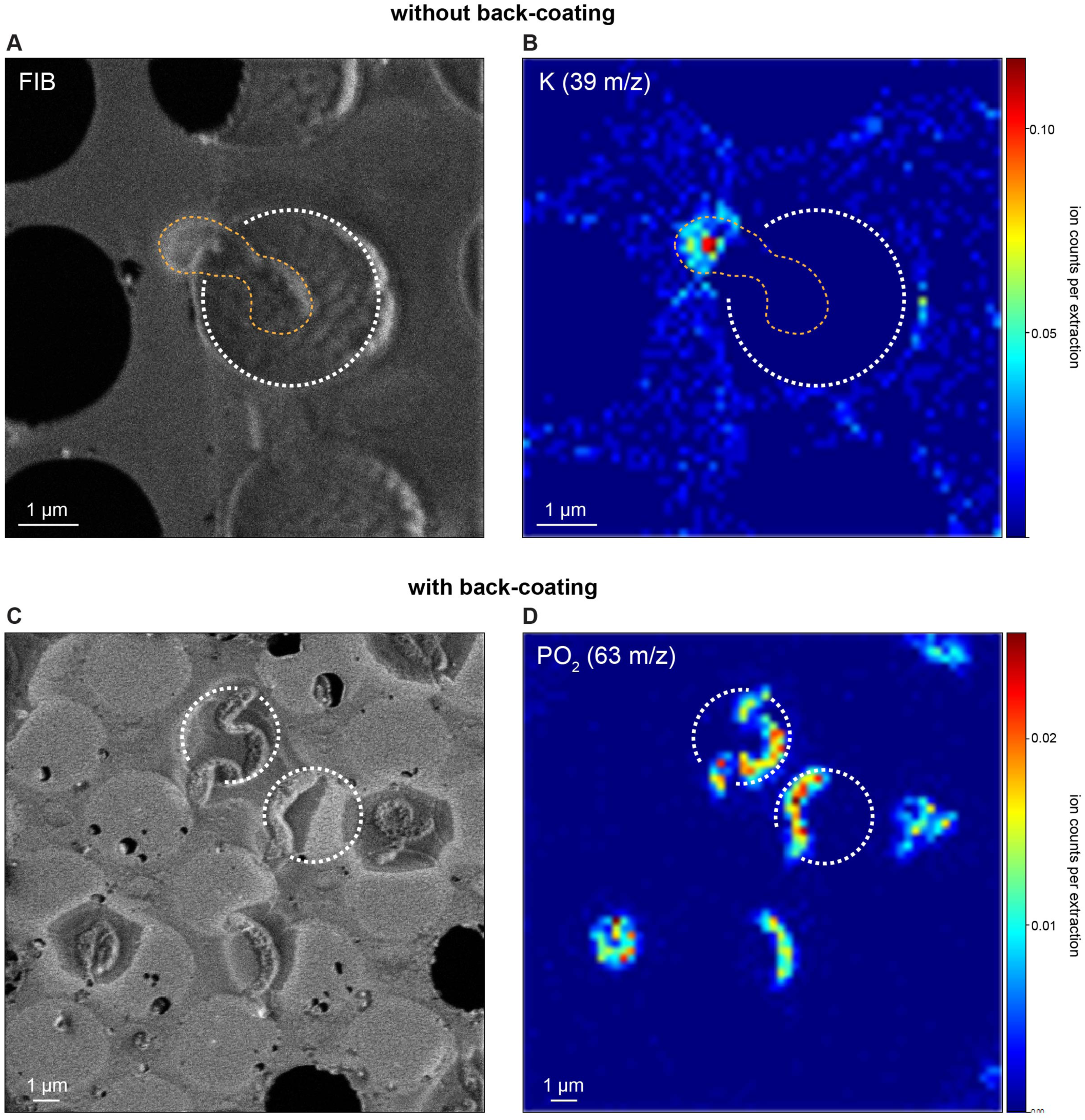
Increase of sample stability and MS signal from holey supports by back-coating. **(A-B)** FIB image **(A)** and cryo-FIB-SIMS distribution of potassium **(B)** in a *C. crescentus* cell on a holey amorphous carbon support without back-coating. In the FIB image, warping of the holes on the left can be observed. Cryo-FIB-SIMS signal is observed from the part of the cell on the carbon support, but not from the cellular area located within the hole in the support foil. **(C-D)** FIB image **(C)** and cryo-FIB-SIMS distribution of PO2 **(D)** of *C. crescentus* cells on holey amorphous carbon support with back-coating. The additional stability due to the organometallic coating reduces warping and facilitates the detection of the cryo-FIB-SIMS signal from the holey areas of the support film.

**Figure S3:**
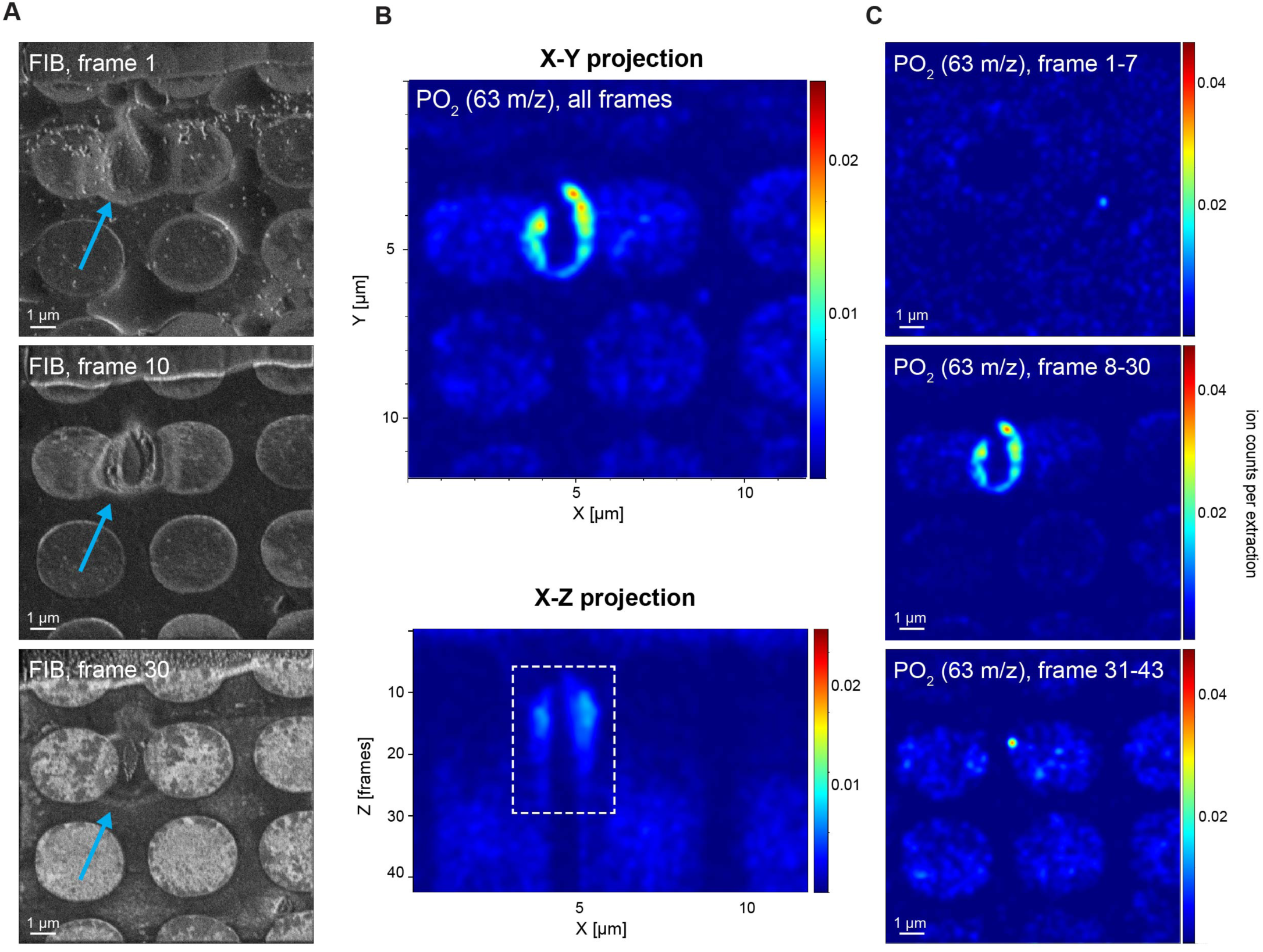
Probing the 3D chemical composition of biological specimens with cryo-FIB-SIMS. **(A)** Secondary electron (FIB) images of the sample corresponding to different frames, i.e. different heights along the ‘Z’-stack. At frame 1 (top), the cell (marked by blue arrow) is still covered in ice. By frame 10 (center), the beam is probing the interior of the cell. At frame 30 (bottom), the cellular material has been fully ablated by the beam. Thus, repeatedly scanning over the area of interest provides 3D information about the sample. **(B)** Corresponding cryo-FIB-SIMS image of the same cell as in **(A)** for 63 *m/z* (PO2) in an X-Y projection of all frames in the stack (top) and an X-Z slice through the Z-stack (bottom). The thickness of the volume shown here (43 frames in total) can be estimated to be between 500-700 nm based on known sputter rates of vitrified ice for Ga beams and the mean thickness of *C. crescentus* cells. While the X-Y projection shows the lateral distribution of the PO2 signal, the X-Z slice yields depth information and allows the localization of the cell along the Z-axis, where cellular signal can be detected from frames 8-30 (white dashed box). This is confirmed by X-Y images resulting from projections of specific frames as shown in **(C)**: the X-Y projection of frames 1-7 (top) does not show cellular signal and corresponds to the ice layer enveloping the cell. The X-Y projection of frames 8-30 shows a cellular image, in accordance with the cellular localization seen in the FIB images and the X-Z slice **(A, B)**. After frame 30 (bottom, frame 31-43), the cell is not visible any more as at that point, it has been fully removed by the interaction with the ion beam (see also Supplementary Video S1).

**Figure S4:**
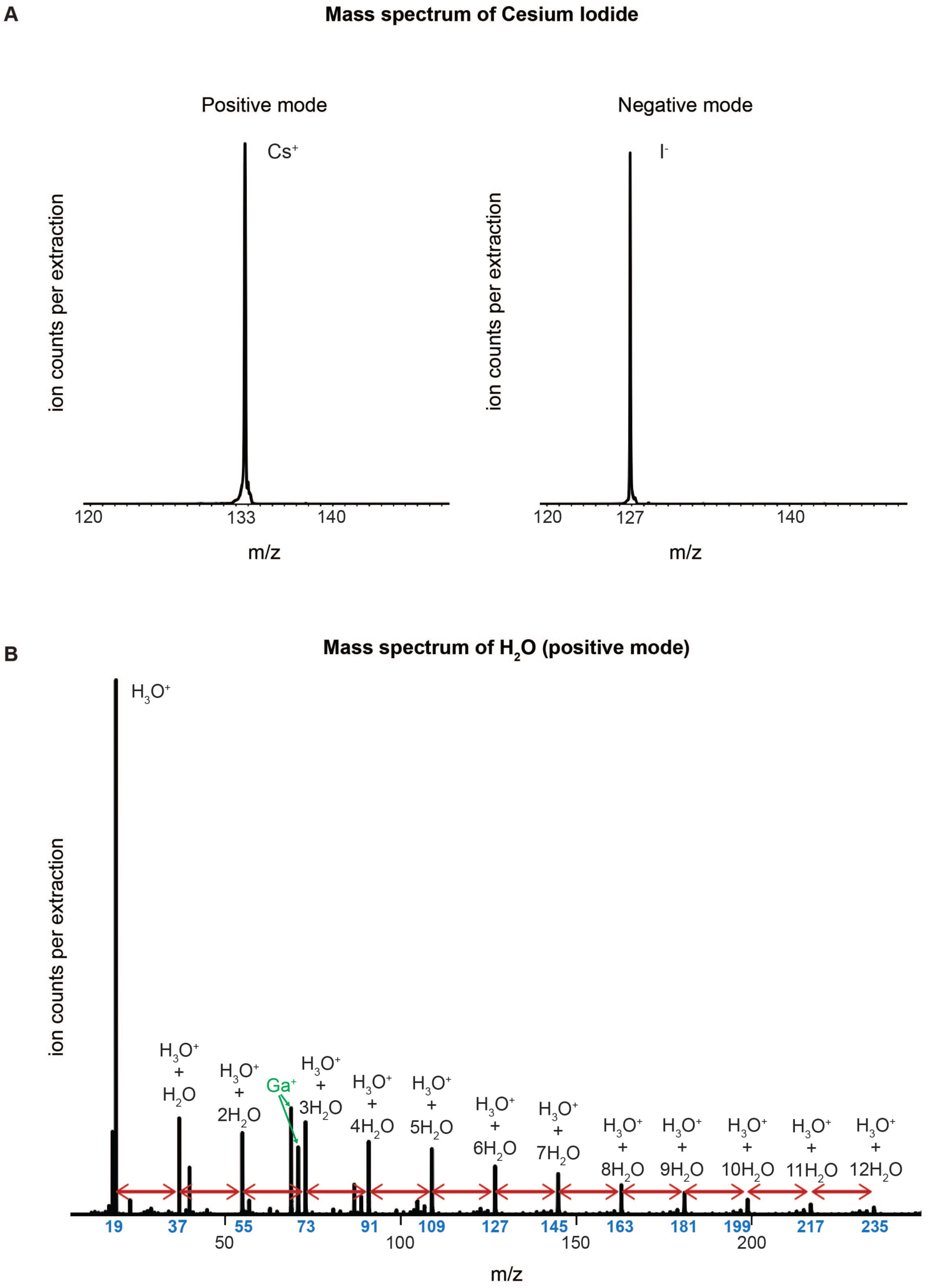
Cryo-EM-FIB-SIMS workflow. **(A)** Main peaks in positive and negative ion mode mass spectra of a vitrified sample of CsI: 133 *m/z*, corresponding to Cs^+^, and 127 *m/z*, corresponding to I^-^ are detected. **(B)** Positive mode mass spectrum of a vitrified sample of H2O showing characteristic water clusters: the main peak is 19 *m/z* (H3O^+^) followed by peaks spaced 18 *m/z* apart, corresponding to clusters with a linearly increasing number of H2O molecules. Peaks due to gallium are highlighted in green.

**Figure S5:**
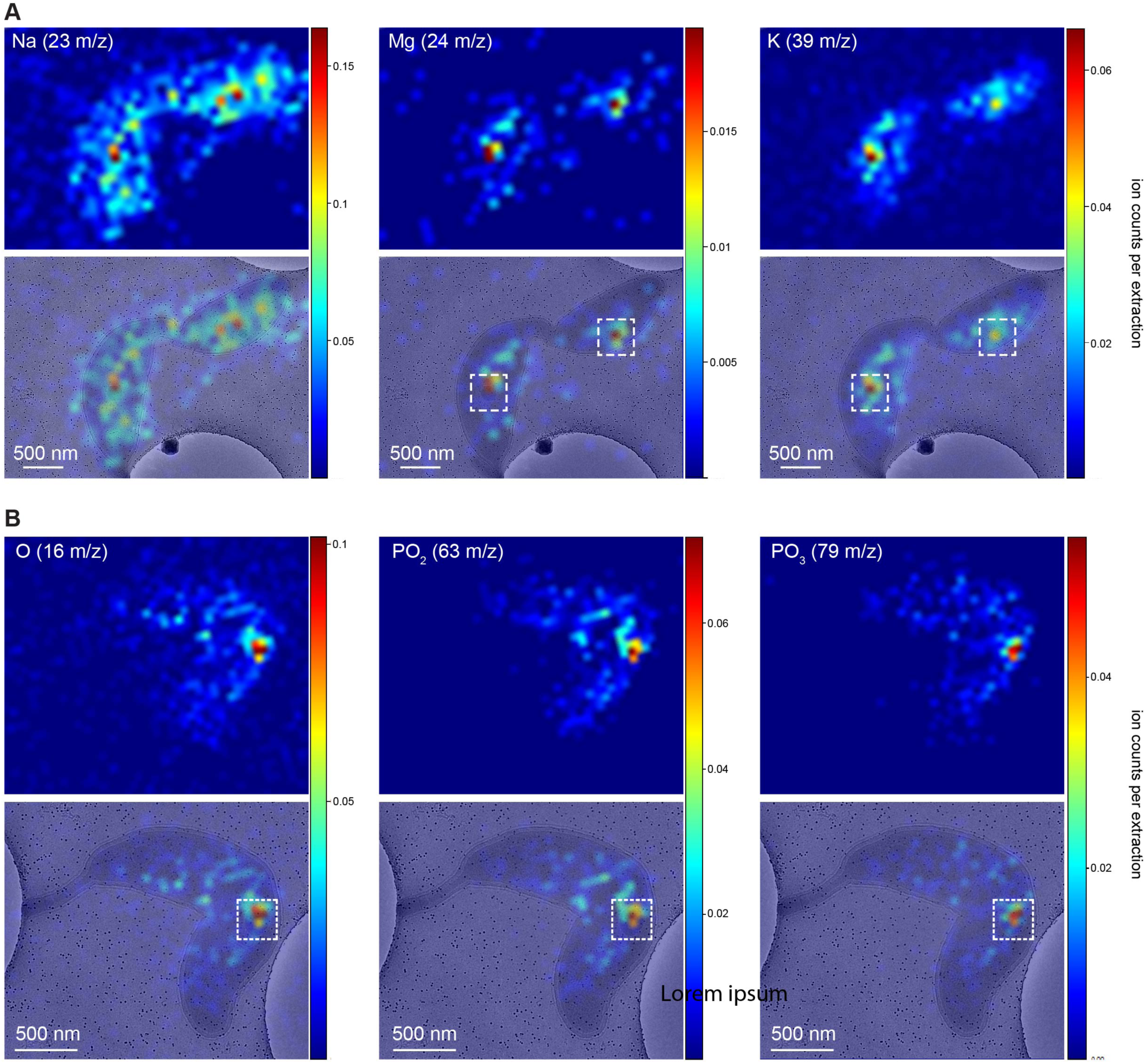
Additional examples demonstrating the cryo-EM-FIB-SIMS workflow on unlabelled *C. crescentus* cells. **(A)** Positive ion mode correlative analysis of a *C. crescentus* cell showing cryo-FIB-SIMS images of three prominent *m/z* peaks (Na, Mg, and K) and the respective overlays with the cryo-EM images. The dashed white boxes indicate locations of storage granules. **(B)** Negative ion mode correlative analysis of a *C. crescentus* cell showing cryo-FIB-SIMS images of three prominent *m/z* peaks (O, PO2, and PO3) and the respective overlays with the cryo-EM images. The dashed white boxes indicate locations of storage granules.

**Figure S6:**
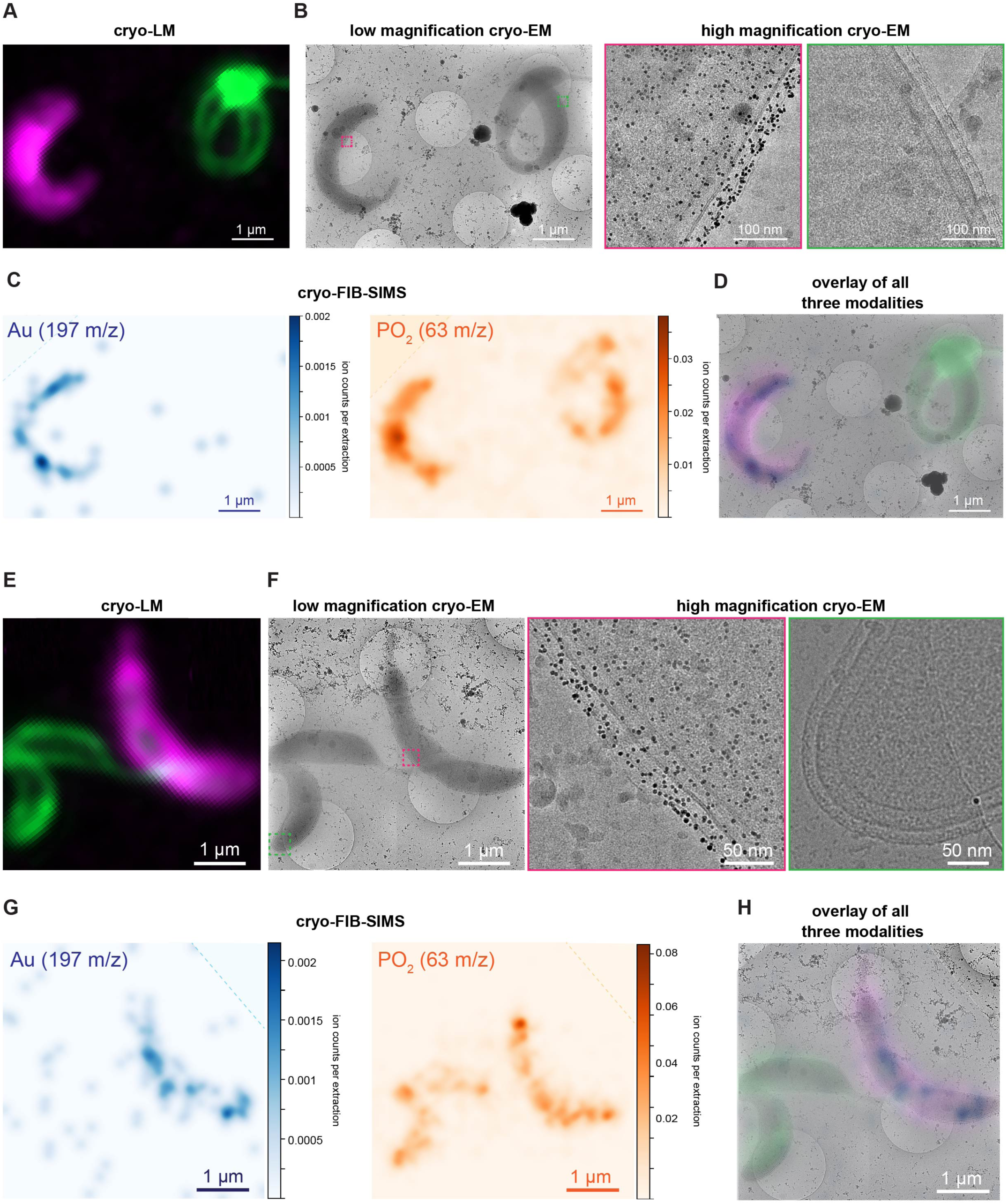
Additional examples demonstrating the cryo-CLEM-FIB-SIMS workflow. **(A, E)** Cryo-LM images showing the dual fluorescent labelling of *C. crescentus* cells. **(B, F)** Imaging the same cells in cryo-EM demonstrates that the magenta-fluorescent cells feature gold nanoparticle labelling (black dots), whereas the green-fluorescent cells do not. **(C, G)** Cryo-FIB-SIMS only detects the magenta+Au cells in the *m/z* channel corresponding to elemental gold (left, blue), while both cells are detected in the PO2 channel (right, orange), which can be further visualized by the overlay of all three imaging modalities **(D, H)**. In **(C)** and **(G)**, there are missing corners of the images due to different sample rotations in the different imaging modalities, which were padded above the dashed lines to match the layout of the cryo-EM and cryo-LM images.

**Figure S7:**
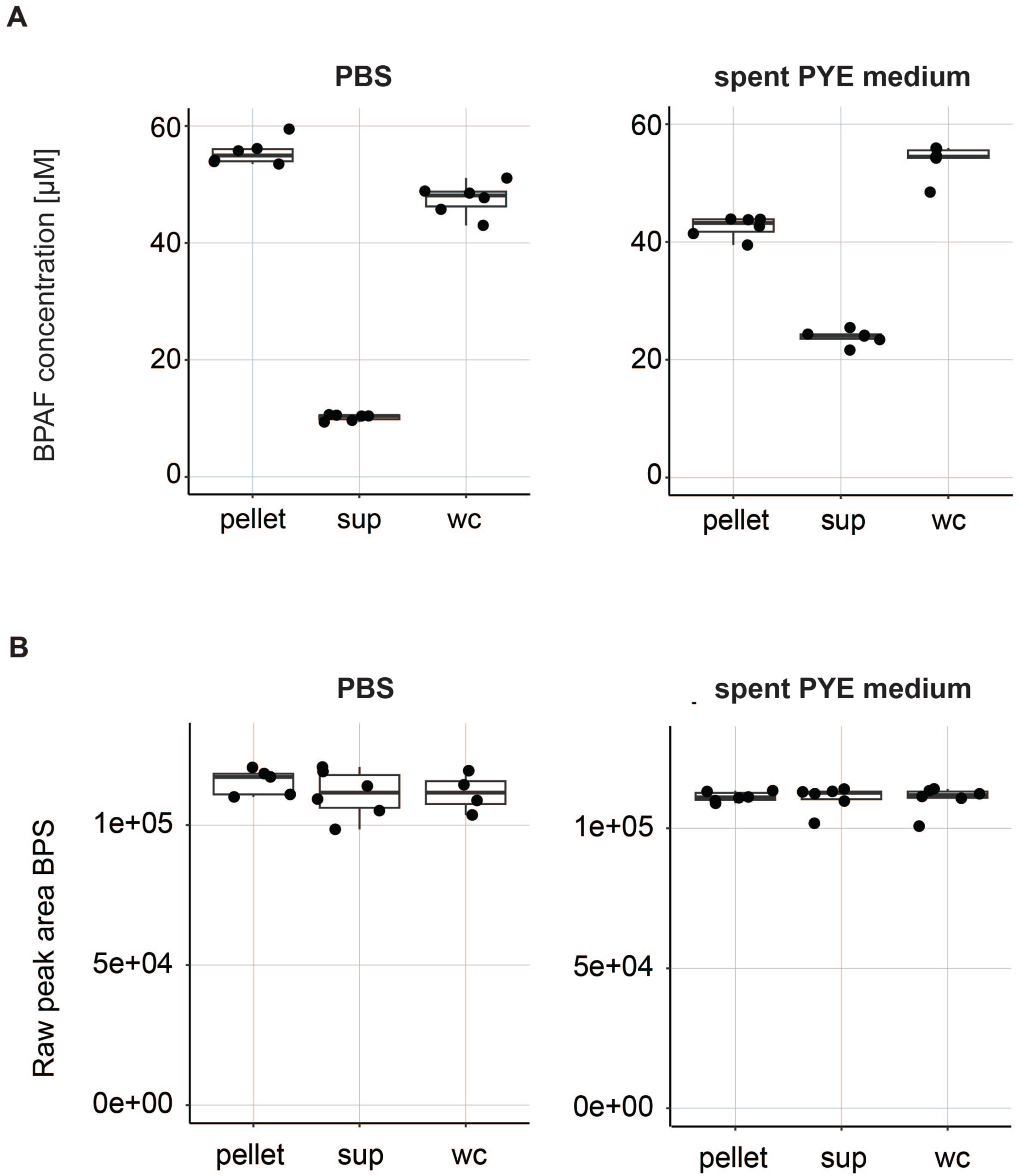
Bioaccumulation of BPAF in *C. crescentus*. Panel **(A)** is repeated here from Fig. 4b to facilitate comparison with the internal standard shown in panel **(B)**. **(A)** Bioaccumulation analysis performed both in PBS and spent PYE (Peptone Yeast Extract) medium shows clear differences in BPAF concentration between supernatant and pellet after exposure to 50 µM BPAF, indicating BPAF accumulation within the cells. wc denotes the whole culture (bacteria plus supernatant, PYE or PBS with final concentrations of 50µM BPAF), sup denotes the supernatant (PYE or PBS with 50 µM BPAF after the removal of the bacteria by spinning), and pellet denotes the cell pellet from the wc (exposed to 50 µM BPAF), reconstituted in spent medium or PBS without BPAF. The higher efficiency of bioaccumulation in PBS compared to the spent PYE medium is expected given the lack of nutrients in PBS. **(B)** The internal standard BPS (bisphenol S) was added after incubation as part of the extraction buffer and shows low variation in sample injection, demonstrating that the pattern observed in a is a biological and not a technical effect.

**Figure S8:**
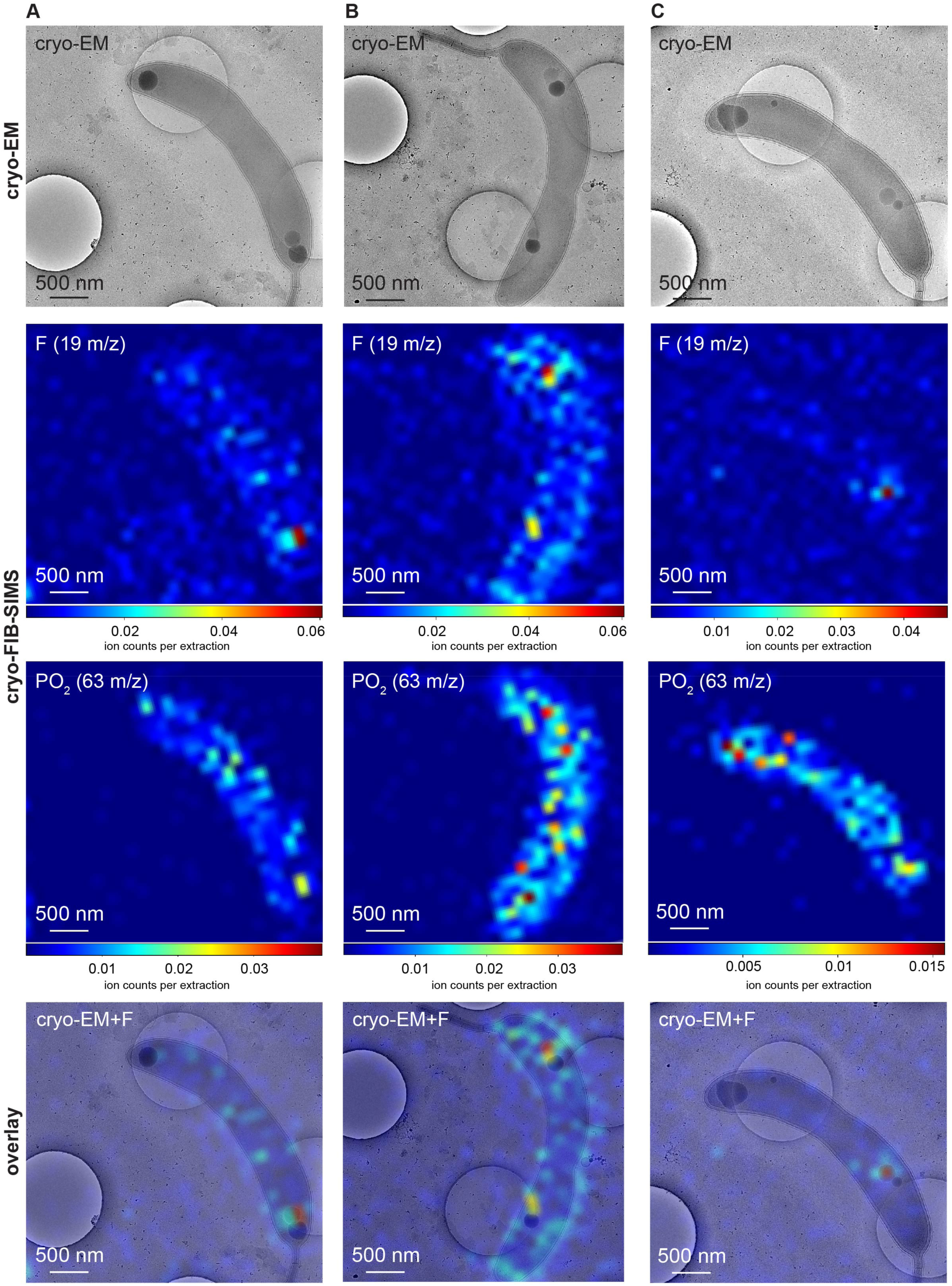
Additional examples demonstrating the uptake of BPAF by *C. crescentus*. **(A-C)** Three different *C. crescentus* cells are shown with BPAF-associated fluorine signal that is co-localised with storage granules after BPAF exposure. Top to bottom: cryo-EM images, cryo-FIB-SIMS images for F and PO2, overlay of the fluorine distributions with the cryo-EM images.

**Figure S9:**
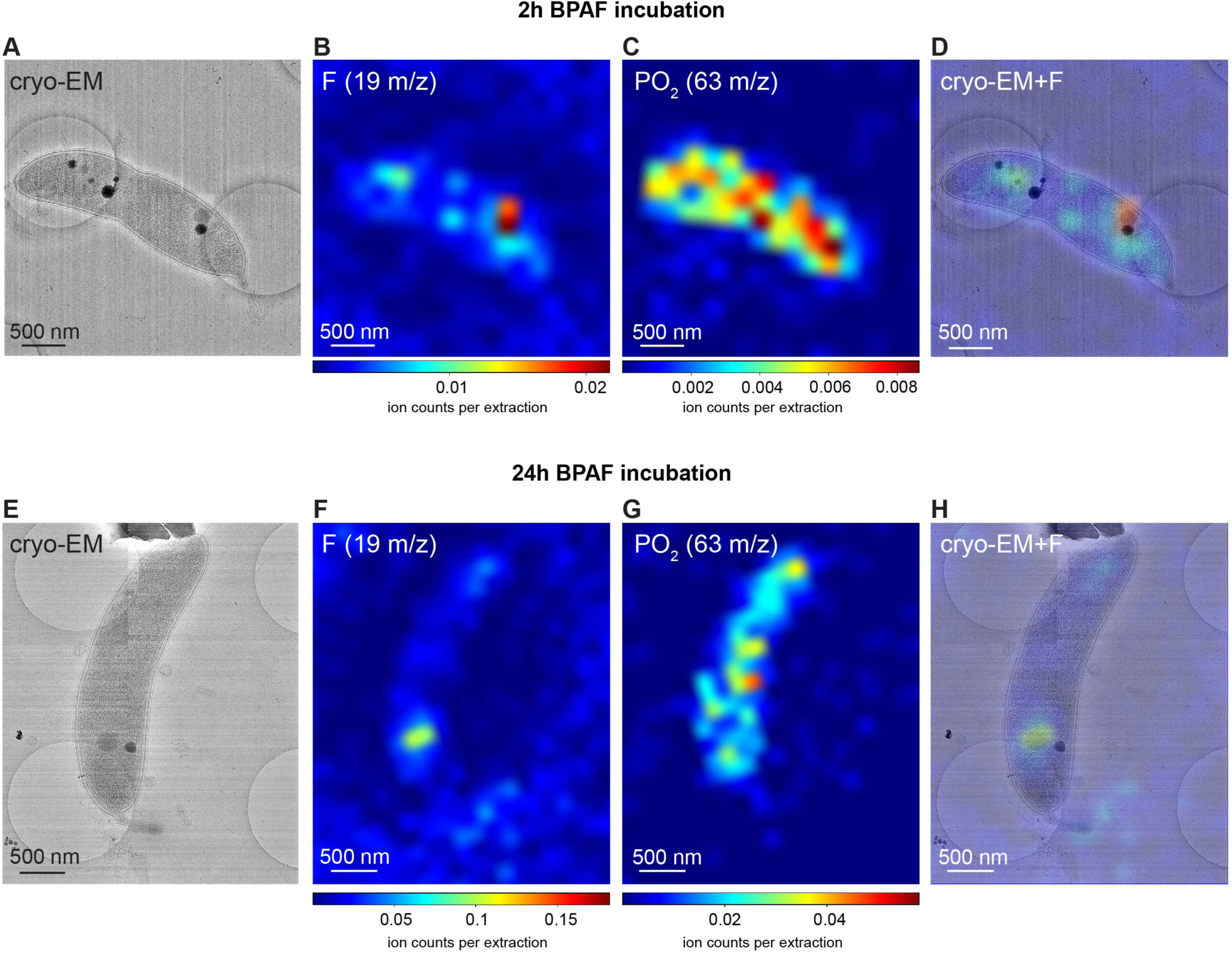
BPAF exposure at different time points. **(A-D)** *C. crescentus* after 2 hours of exposure to 100 μM BPAF: cryo-EM image **(A)**, cryo-FIBSIMS images for fluorine **(B)** and PO2 **(C)**, and overlay of the fluorine distribution with the cryo-EM image **(D)** show that already after 2 hours, there is uptake and localization of BPAF in specific intracellular locations. **(E-H)** *C. crescentus* after 24 hours of exposure to 100 μM BPAF: cryo-EM image **(E)**, cryo-FIB-SIMS images for F **(F)** and PO2 **(G)**, and overlay of the fluorine distribution with the cryo-EM image **(H)** show that BPAF remains within the cells with signal localized to the storage granules in the cytosol.

## References

1. Beck, M. & Baumeister, W. Cryo-Electron Tomography: Can it Reveal the Molecular Sociology of Cells in Atomic Detail? Trends in Cell Biology 26, 825–837 (2016).

2. Henderson, R. The potential and limitations of neutrons, electrons and X-rays for atomic resolution microscopy of unstained biological molecules. Quart. Rev. Biophys. 28, 171–193 (1995).

3. Ochner, H. & Bharat, T. A. M. Charting the molecular landscape of the cell. Structure 31, 1297–1305 (2023).

4. Pillatsch, L., Östlund, F. & Michler, J. FIBSIMS: A review of secondary ion mass spectrometry for analytical dual beam focussed ion beam instruments. Prog. Cryst. Growth Charact. Mater. 65, 1–19 (2019).

5. Stevie, F. A. Focused Ion Beam Secondary Ion Mass Spectrometry (FIB-SIMS). in Introduction to Focused Ion Beams (eds. Giannuzzi, L. A. & Stevie, F. A.) 269–280 (Springer US, Boston, MA, 2005). doi:10.1007/0-387-23313-X_13.

6. Winograd, N. Imaging Mass Spectrometry on the Nanoscale with Cluster Ion Beams. Anal. Chem. 87, 328–333 (2015).

7. Fletcher, J. S. Cellular imaging with secondary ion mass spectrometry. Analyst 134, 2204 (2009).

8. McDonnell, L. A. & Heeren, R. M. A. Imaging mass spectrometry. Mass Spectrom. Rev. 26, 606–643 (2007).

9. Passarelli, M. K. et al. The 3D OrbiSIMS—label-free metabolic imaging with subcellular lateral resolution and high mass-resolving power. Nat Methods 14, 1175–1183 (2017).

10. Kotowska, A. M. et al. Protein identification by 3D OrbiSIMS to facilitate in situ imaging and depth profiling. Nat. Commun. 11, 5832 (2020).

11. Tian, H., Six, D. A., Krucker, T., Leeds, J. A. & Winograd, N. Subcellular Chemical Imaging of Antibiotics in Single Bacteria Using C60 -Secondary Ion Mass Spectrometry. Anal. Chem. 89, 5050–5057 (2017).

12. Tian, H. et al. Successive High-Resolution (H2O)n-GCIB and C60-SIMS Imaging Integrates Multi-Omics in Different Cell Types in Breast Cancer Tissue. Anal. Chem. 93, 8143–8151 (2021).

13. Zhang, J. et al. Cryo-OrbiSIMS for 3D Molecular Imaging of a Bacterial Biofilm in Its Native State. Anal. Chem. 92, 9008–9015 (2020).

14. Gilmore, I. S., Heiles, S. & Pieterse, C. L. Metabolic Imaging at the Single-Cell Scale: Recent Advances in Mass Spectrometry Imaging. *Annual Rev*. Anal. Chem. 12, 201–224 (2019).

15. De Castro, O. et al. npSCOPE: A New Multimodal Instrument for In Situ Correlative Analysis of Nanoparticles. Anal. Chem. 93, 14417–14424 (2021).

16. Lindell, A. E. et al. Extensive PFAS accumulation by human gut bacteria. Preprint at 10.1101/2024.09.17.613493 (2024).

17. von Kügelgen, A., et al. *In Situ* Structure of an Intact Lipopolysaccharide-Bound Bacterial Surface Layer. Cell 180, 348–358.e15 (2020).

18. Herdman, M. et al. Cell cycle dependent coordination of surface layer biogenesis in *Caulobacter crescentus*. Nat. Commun. 15, 3355 (2024).

19. De Koning, E. A. et al. The PHB Granule Biogenesis Pathway in *Caulobacter*. Preprint at 10.1101/2023.07.06.548030 (2023).

20. Henry, J. T. & Crosson, S. Chromosome replication and segregation govern the biogenesis and inheritance of inorganic polyphosphate granules. Mol. Biol. Cell. 24, 3177–3186 (2013).

21. Boutte, C. C., Henry, J. T. & Crosson, S. ppGpp and Polyphosphate Modulate Cell Cycle Progression in *Caulobacter crescentus*. J. Bacteriol. 194, 28–35 (2012).

22. Comolli, L. R., Kundmann, M. & Downing, K. H. Characterization of intact subcellular bodies in whole bacteria by cryo-electron tomography and spectroscopic imaging. J. Microsc. 223, 40–52 (2006).

23. Kukulski, W. et al. Correlated fluorescence and 3D electron microscopy with high sensitivity and spatial precision. J. Cell Biol. 192, 111–119 (2011).

24. Moser, F. et al. Cryo-SOFI enabling low-dose super-resolution correlative light and electron cryo-microscopy. Proc. Natl. Acad. Sci. U.S.A. 116, 4804–4809 (2019).

25. Herdman, M. et al. High-resolution mapping of metal ions reveals principles of surface layer assembly in *Caulobacter crescentus* cells. Structure 30, 215–228.e5 (2022).

26. Caspy, I., Wang, Z. & Bharat, T. A. M. Structural biology inside multicellular specimens using electron cryotomography. Quart. Rev. Biophys. 58, e6 (2025).

27. Marko, M., Hsieh, C., Schalek, R., Frank, J. & Mannella, C. Focused-ion-beam thinning of frozen-hydrated biological specimens for cryo-electron microscopy. Nat. Methods 4, 215–217 (2007).

28. Heymann, J. A. W. et al. Site-specific 3D imaging of cells and tissues with a dual beam microscope. J. Struct. Biol. 155, 63–73 (2006).

29. Baumeister, W. Electron tomography: towards visualizing the molecular organization of the cytoplasm. Current Opinion in Structural Biology 12, 679–684 (2002).

30. Escrivá, L., Hanberg, A., Zilliacus, J. & Beronius, A. Assessment of the endocrine disrupting properties of Bisphenol AF according to the EU criteria and ECHA/EFSA guidance. EF*S2* 17, (2019).

31. Priebe, A. & Michler, J. Review of Recent Advances in Gas-Assisted Focused Ion Beam Time-of-Flight Secondary Ion Mass Spectrometry (FIB-TOF-SIMS). Materials 16, 2090 (2023).

32. Winograd, N. The Magic of Cluster SIMS. Anal. Chem. 77, 142 A-149 A (2005).

33. Lange, F. et al. Correlative fluorescence microscopy, transmission electron microscopy and secondary ion mass spectrometry (CLEM-SIMS) for cellular imaging. PLoS ONE 16, e0240768 (2021).

34. Lork, A. A. et al. Subcellular protein turnover in human neural progenitor cells revealed by correlative electron microscopy and nanoscale secondary ion mass spectrometry imaging. Chem. Sci. 15, 3311–3322 (2024).

35. Rabasco, S. et al. Characterization of Stress Granule Protein Turnover in Neuronal Progenitor Cells Using Correlative STED and NanoSIMS Imaging. Int. J. Mol. Sci. 24, 2546 (2023).

36. Esser, T. K. et al. Cryo-EM of soft-landed β-galactosidase: Gas-phase and native structures are remarkably similar. Sci. Adv. 10, eadl4628 (2024).

37. O’Reilly, F. J. et al. In-cell architecture of an actively transcribing-translating expressome. Science 369, 554–557 (2020).

38. Ramakrishna, P. et al. Elemental cryo-imaging reveals SOS1-dependent vacuolar sodium accumulation. Nature 637, 1228–1233 (2025).

39. Pfeil-Gardiner, O. et al. Elemental mapping in single-particle reconstructions by reconstructed electron energy-loss analysis. Nat Methods (2024) doi:10.1038/s41592-024-02482-5.

40. Bharat, T. A. M. et al. Structure of the hexagonal surface layer on Caulobacter crescentus cells. Nat Microbiol 2, 17059 (2017).

41. Sulkowski, N. I., Hardy, G. G., Brun, Y. V. & Bharat, T. A. M. A Multiprotein Complex Anchors Adhesive Holdfast at the Outer Membrane of Caulobacter crescentus. J Bacteriol 201, e00112–19 (2019).

42. Herdman, M. et al. Cell cycle dependent coordination of surface layer biogenesis in Caulobacter crescentus. Nat Commun 15, 3355 (2024).

43. Herdman, M. et al. High-resolution mapping of metal ions reveals principles of surface layer assembly in Caulobacter crescentus cells. Structure 30, 215–228.e5 (2022).

44. Isbilir, B., Yeates, A., Alva, V. & Bharat, T. A. M. Mapping the ultrastructural topology of the corynebacterial cell surface. 2024.09.05.611374 Preprint at 10.1101/2024.09.05.611374 (2024).

45. Kelley, K. et al. Waffle Method: A general and flexible approach for improving throughput in FIB-milling. Nat Commun 13, 1857 (2022).

46. Fu, J., Joshi, S. B. & Catchmark, J. M. Sputtering rate of micromilling on water ice with focused ion beam in a cryogenic environment. Journal of Vacuum Science & Technology A: Vacuum, Surfaces, and Films 26, 422–429 (2008).

47. Schindelin, J., et al. Fiji: an open-source platform for biological-image analysis. Nat Methods 9, 676–682 (2012).

48. Mastronarde, D. N. Automated electron microscope tomography using robust prediction of specimen movements. Journal of Structural Biology 152, 36–51 (2005).

49. Lowe, D. G. Distinctive Image Features from Scale-Invariant Keypoints. International Journal of Computer Vision 60, 91–110 (2004).

50. Zheng, S., et al. AreTomo: An integrated software package for automated marker-free, motion-corrected cryo-electron tomographic alignment and reconstruction. Journal of Structural Biology: X 6, 100068 (2022).

51. Kremer, J. R., Mastronarde, D. N. & McIntosh, J. R. Computer Visualization of Three-Dimensional Image Data Using IMOD. Journal of Structural Biology 116, 71–76 (1996).

52. Laurent Gatto, Christophe Vanderaa. QFeatures. Bioconductor 10.18129/B9.BIOC.QFEATURES.

53. Smith, Tom. biomasslmb: BioMass LMB Proteomics Data Analysis Utility Functions. R package version 0.0.1. (2025).

54. Plubell, D. L. et al. Extended Multiplexing of Tandem Mass Tags (TMT) Labeling Reveals Age and High Fat Diet Specific Proteome Changes in Mouse Epididymal Adipose Tissue. Molecular & Cellular Proteomics 16, 873–890 (2017).

55. Ritchie, M. E. et al. limma powers differential expression analyses for RNA-sequencing and microarray studies. Nucleic Acids Research 43, e47–e47 (2015).

56. Yoav Benjamini & Yosef Hochberg. Controlling the False Discovery Rate: A Practical and Powerful Approach to Multiple Testing. Journal of the Royal Statistical Society. Series B (Methodological) 57, 289–300 (1995).

